# Unravelling the role of oceanographic connectivity in intra-specific diversity of marine forests at global scale

**DOI:** 10.1101/2023.07.12.548579

**Authors:** Térence Legrand, Eliza Fragkopoulou, Lauren Vapillon, Lidiane Gouvêa, Ester A. Serrão, Jorge Assis

## Abstract

**Aim:** Intra-specific diversity results from complex interactions of intermingled eco-evolutionary processes along species’ history, but their relative contribution has not been addressed at the global scale. Here, we unravel the role of present-day oceanographic connectivity in explaining the genetic differentiation of marine forests across the ocean.

**Location:** Global.

**Time period:** Contemporary.

**Major taxa studied:** Marine forests of brown macroalgae (order Fucales, Ishigeales, Laminariales, Tilopteridale).

**Methods:** Through systematic literature revision, we compiled a comprehensive dataset of genetic differentiation, encompassing 662 populations of 34 species. A biophysical model coupled with network analyses estimated multigenerational oceanographic connectivity and centrality across the marine forest global distribution. This approach integrated species’ dispersive capacity and long-distance dispersal events. Linear mixed models tested the relative contribution of site-specific processes, connectivity, and centrality in explaining genetic differentiation.

**Results:** We show that spatiality dependent eco-evolutionary processes, as described by our models, are prominent drivers of genetic differentiation in marine forests (significant models in 92.6 % of the cases with an average R^2^ of 0.49 ± 0.07). Specifically, we reveal that 19.6 % of variance is explicitly induced by contemporary connectivity and centrality. Moreover, we demonstrate that LDD is key in connecting populations of species distributed across large water masses and continents.

**Main conclusions:** We deciphered the role of present-day connectivity in observed patterns of genetic differentiation of marine forests. Our findings significantly contribute to the understanding of the drivers of intra-specific diversity on a global scale, with implications for biogeography and evolution. These results can guide well-informed conservation efforts, including the designation of marine protected areas, as well as spatial planning for genetic diversity in aquaculture, which is particularly relevant for sessile ecosystems structuring species such as brown macroalgae.

## Introduction

Intra-specific diversity is critical for ecosystem resilience, species adaptation and supporting nature’s contribution to people (Des Roches et al., 2018, Des Roches et al., 2021, Hoban et al., 2020). Over the past decades, a wide range of genotypic variation among populations has been highlighted by molecular genetics and genomic tools, particularly in marine species (Selkoe et al., 2016). The contrasting observed patterns, ranging between highly structured and panmictic populations, result from complex interactions of eco-evolutionary processes throughout the species’ history (Lowe et al., 2017). Among these processes, gene flow promoted by population connectivity through dispersal has a pivotal multifaceted impact on genetic differentiation across various time scales (Lowe & Allendorf, 2010). Indeed, observed genetic structures can be attributed in part to past gene flow patterns, with historical barriers promoting long-term differentiation (e.g., Arnaud-Haond et al., 2007) and colonisation events facilitating the spread of low-frequency alleles at the front of species’ ranges (Waters et al., 2013). On top of that, contemporary gene flow between established populations can homogenise allele frequencies, reshuffle mutations, disperse adaptive changes, and mitigate the effects of inbreeding depression (Hellberg, 2009, Lenormand et al., 2002). However, deciphering the relative contribution of present-day connectivity from historical processes on observed genetic structures remains a major challenge, with attempts at the global scale limited to a few species (e.g., mangrove forests; Gouvêa et al., 2023) and regions (e.g., Mediterranean Sea; Legrand et al., 2022). Addressing this knowledge gap is not only crucial for biogeography but also for biodiversity conservation. This can identify key areas for preserving unique genetic diversity (Andrello et al., 2022), guide the designation of Marine Protected Areas (Assis et al., 2021, Abecassis et al., 2023), and inform aquaculture spatial planning (Brackel et al., 2021), especially relevant for sessile ecosystems structuring species like marine forests that rely on oceanic currents for dispersal.

Marine forests of large brown algae play a vital role in coastal areas, ranked among the most productive and biodiversity-rich ecosystems worldwide. These blue-green canopy-forming macroalgae provide essential ecological services, including carbon sequestration, nutrient cycling, and coastal protection (e.g., sediment stabilisation and protection against ocean waves). They further support critical habitats for numerous associated species, many of which are of high commercial value (Eger et al., 2023; Araújo et al., 2016; Christie et al., 2009) and a growing aquaculture industry, whether for direct human consumption or climate change mitigation (Gouvêa et al., 2020). Marine forests have restricted gamete, zygote and/or spore dispersal (here referred to as “in-house” dispersal) and therefore tend to display fine-scale genetic structure with high levels of genetic endemicity across species’ distribution ranges (Assis et al., 2018, 2023, Maggs et al., 2008). Although rare, long-distance dispersal (LDD) events can also occur through detached fragments or thalli (here referred to as “third-party” dispersal), which may remain viable and buoyant for long periods (Thiel & Gutow, 2005). These events extend the dispersal capacity of marine forests, facilitating the successful establishment of populations across large water masses spanning hundreds of kilometres (Bernardes Batista et al., 2018; Assis et al., 2023). The interplay between frequent in-house and rare LDD events determines the scales at which marine forests’ populations are connected and exchange genes. Most studies focused on the genetic drivers of marine forests have inferred the contribution of present-day connectivity through observed patterns of population genetic differentiation (Maggs et al., 2008) or through isolation by distance models (Riquet et al., 2021). These simplistic approaches cannot capture the complexity of ongoing oceanographic processes (Jahnke et al., 2022), which can produce asymmetrical population gene flow with high temporal and spatial variability (Gouvêa et al., 2023, Legrand et al., 2022). The few studies considering mechanistic biophysical modelling mimicking oceanographic processes over multiple generations are limited to few species (e.g., *Cystoseira amentacea, Fucus radicans, Fucus vesiculosus, Laminaria ochroleuca* and *Laminaria pallida*) and realms (e.g., Temperate Northern Atlantic and Temperate Southern Africa; Spalding et al., 2007) and have never integrated LDD (Assis et al., 2018, 2022, Buonomo et al., 2017, Pereyra et al., 2013), precluding conclusions on the relative contribution of present-day oceanographic connectivity on marine forests’ genetic structures.

Here, we investigate the relative contribution of present-day oceanographic connectivity on the intra-specific diversity of marine forests. We developed a biophysical model that estimated oceanographic connectivity across the global distribution of marine forests, while considering their dispersive capacity and sporadic LDD. Simulations were coupled with network analysis to account for multigenerational stepping-stone connectivity and centrality, and results were tested against empirical genetic data, collected through a systematic literature revision. Linear mixed models disentangled the role of oceanographic connectivity and centrality from other spatial processes in explaining the genetic differentiation of marine forests. Our findings have important implications for marine forests’ biogeography and evolution, as well as for conservation and management, including the designation of marine protected areas (MPAs), aquaculture spatial planning, and climate change research exploring future impacts on gene flow.

## Methods

### Marine forest genetic differentiation dataset

We conducted a systematic search in the Web of Science platform (accessed in October 2022) to compile genetic differentiation data for marine forests of brown macroalgae spanning the last three decades. We used the combination of the words “genetic diversity” or “genetic differentiation” or “phylogeography” with “algae” or “macroalgae” or “seaweed” or “heterokontophyta”. We selected studies that provided pairwise population genetic differentiation estimations. For each chosen study, we extracted information regarding the number of populations and sampled individuals, the geographic location, the molecular marker utilised in the genetic analyses, the differentiation index and the pairwise matrix of genetic differentiation (SI Table S2). We reassessed the selected studies based on updated taxonomic and ecological information. Notably, we divided the sampled populations of *Undaria pinnatifida* (Shan et al., 2019) into two distinct datasets, representing the native Chinese and non-native European populations. Similarly, we splitted *Macrocystis pyrifera* populations into datasets based on the northern and southern hemispheres (Assis et al., 2023). Regarding the two studies investigating the impact of marine heatwaves on genetic differentiation (Coleman et al., 2020; Gurgel et al., 2020), we treated the ‘before’, ‘after’, and ‘before vs. after’ time points as separate datasets. It is important to note that *Halidrys dioica* (Lu et al., 1994) has been updated to its accepted name, *Stephanocystis dioica*.

### Biophysical modelling

We generated simulations of oceanographic connectivity with a biophysical modelling framework widely implemented elsewhere and validated against genetic data of different ecological groups (e.g., macroalgae in Buonomo et al., 2017, Assis et al., 2018, 2022 and mangroves in Gouvêa et al., 2023). This framework tracks passive Lagrangian particles by integrating daily gridded velocity fields provided by the Copernicus Marine Environment Monitoring Service (CMEMS, https://doi.org/10.48670/moi-00021), a data-assimilative operational ocean model implemented on the global ocean at 1/12° horizontal resolution (approximately 8 km). Connectivity simulations were conducted over the global marine distribution (Figure 1), as inferred by stacked species distribution model (SDM) provided by Fragkopoulou et al., 2022. These models provided distribution estimates for nearly all the species of our dataset except for *Fucus guiryi*, *Ishige okamurae*, *Nereia lophocladia*, and *Pylaiella littoralis*. As the observed distributions of these four species (according to AlgaeBase, Guiry & Guiry, 2023) overlapped with our computed global marine forest distribution, we consider the latter as a comprehensive representation of our compiled species.

**Figure 1:**
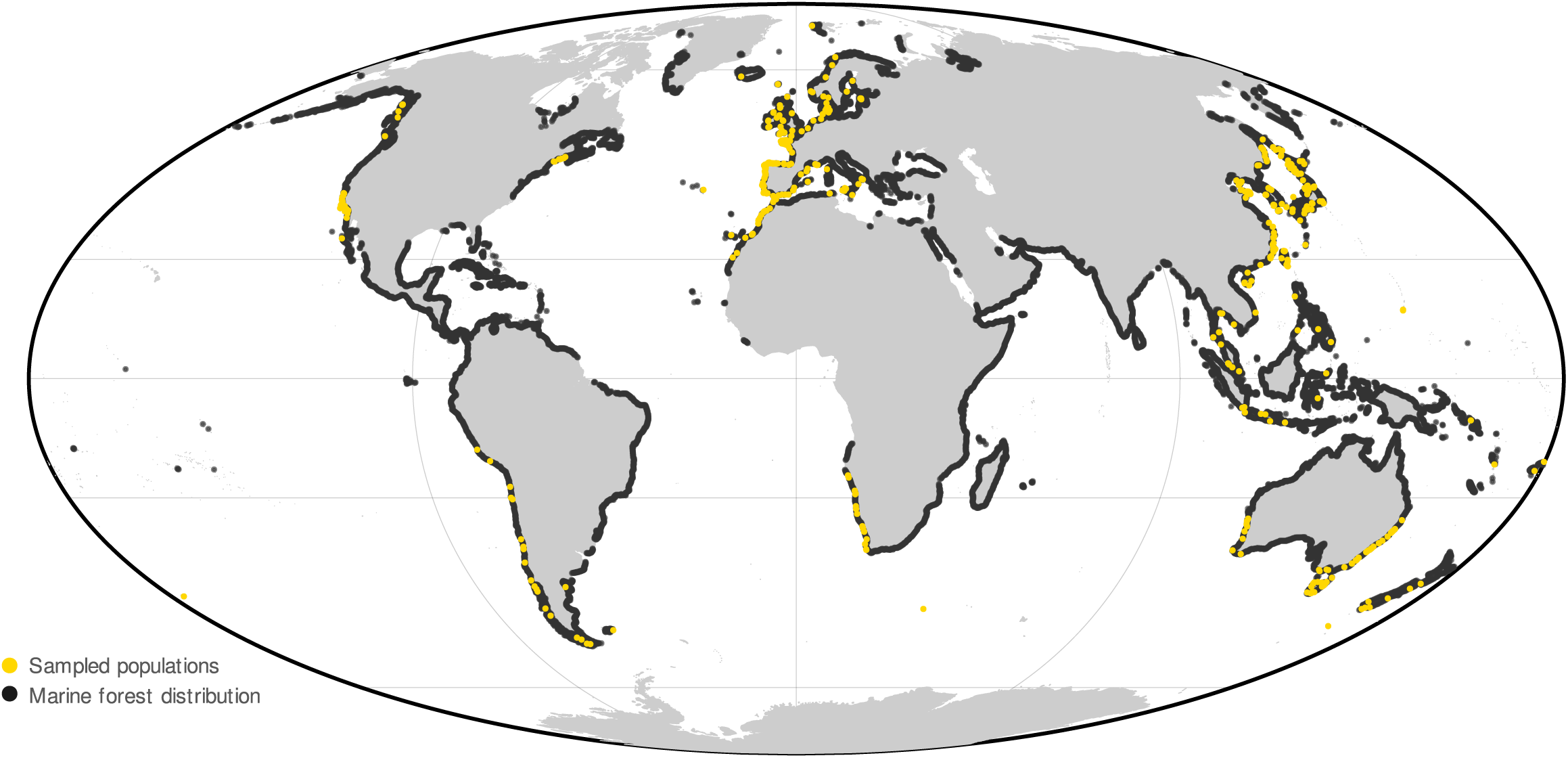
Global distribution of marine forest (dark contour) over which the biophysical model simulated oceanographic connectivity and sampled populations for genetic differentiation data (yellow dots).

The biophysical model released individual particles daily from the centre of sites (hereafter referred to as source and sink sites) with 8.45 km hexagons edge, implemented along the coastline over the global marine forest distribution (n = 16489 source/sink sites, Figure 1). The particles were released over the 11-year period (2010-2020) and advected upon the ocean surface (top vertical layer of the CMEMS model) for up to 180 days until they eventually ended up on sink sites or got lost in the open ocean. The position of each particle was calculated hourly with a bilinear interpolation following the Courant–Friedrichs–Lewy condition. We assign for each sink site, the number of particles incoming for source sink, and the associated mean trajectory time in hours.

### Marine forest dispersal capacity

We gathered data about macroalgae dispersal capacity by combining (i) information on potential dispersal capacity reported on the genetic differentiation studies and (ii) conducting a systematic review on the Web of Science platform (accessed in May 2023) using “dispersal” as a topic, refine by “macroalgae” and “days” or “weeks” or “months”. This step also included Rhodophyta and Chlorophyta species. We selected studies reporting laboratory/field experiments/observations of the potential duration of dispersal (e.g., reproductive viability of fragment, raft survival duration, spore settlement rate, see SI Table S1). Our dataset highlights the different dispersal strategies used by macroalgae introduced earlier: from in-house dispersal by sexual way over a short period of time (gametes, zygotes, and/or spore release) to third-party dispersal by detached algae’s viable fragment or rafts promoting LDD. Because information was scarce and did not cover all species comprised in the genetic dataset (SI Table S2), we summarised information on dispersal duration with a normal distribution fitted on in-house dispersal capacity (μ = mean of dispersal range time, 7.43 d, and σ = standard deviation of dispersal range time, 7.89 d), and normalise the probability density function obtained into a weighting factor function ranging from 0 to 1 (see Figure 4a, following the approach in Di Stefano et al., 2023). We considered LDD events starting at 33 days (i.e., maximal in-house dispersal capacity, Deysher et al., 1981) with a unique weighting factor of 0.01, which corresponds to the proportion of LDD events observed using population assignment tests (D’Aloia et al., 2022). We later weighted, for each sink site, the number of particles incoming for source site by the weighting factor according to the mean trajectory time.

### Multigeneration connectivity and centrality using graph theory

We produced a unique matrix of pairwise probability of connectivity by dividing the weighted number of particles received in sink site from source site by the total weighted number of particles received in sink site, applying the “backward-in-time” dispersal probability approach in Legrand et al. (2022). This approach emphasizes the immigration process, which is instinctively more consistent with genetic theory (i.e., at time t, newly arriving genes at t - 1 are more important than exported genes at t + 1). We obtained a square 16489 x 16489 matrix, summarising the probability of connectivity among different sink and source sites across the global marine forest distribution and considering the weighting function that integrates their diverse dispersal capacities. To evaluate the impact of such functions, we also generate an additional square matrix using a fixed dispersal time of 7.43 d (i.e., the mean of dispersal range time for in-house dispersal capacity). This matrix only considered trajectories with a dispersal duration equal to or less than this fixed time, without any weighting process, as commonly implemented elsewhere (e.g., Gouvêa et al., 2023, Legrand et al., 2019, 2022).

We used graph theory to compute asymmetrical multigeneration connectivity between all sampling localities (i.e., the closest source and sink sites) described in all studies comprised by our genetic differentiation dataset. We set a graph of 16489 nodes representing the sites, while the edges are indexed by the probabilities of connectivity. We use the Dijkstra algorithm to find the shortest path between sampling sites by minimising the sum of distance-transformed probabilities (i.e., - log(probabilities); Dijkstra, 1959). We later determined multi-generation connectivity by multiplying edge probabilities along the inferred shortest paths. We counted the number of stepping-stones needed to connect sampling sites (i.e., A -> B is 0 step, and A -> X -> B is 1 step). To match the symmetric feature of genetic differentiation, we transformed the anisotropic multigeneration probability between pairs of sampling sites to a symmetric probability with P = P_AB_ + P_BA_ - P_AB_P_BA_ (Legrand et al., 2022). Note that we computed the average pairwise genetic differentiation levels when two or more sampled populations were found within a site (Gouvêa et al., 2023). In addition, we utilised a centrality graph-based index to assess the potential role of centrality in population connectivity (Assis et al., 2021; Gouvêa et al., 2023). We considered the harmonic centrality metric, which calculates, for each node, the inverse of the total length of the shortest paths between that node and all other nodes (Marchiori & Latora, 2000). This metric is similar to the closeness centrality (Gouvêa et al., 2023), but it accommodates non-connected graphs, which is applicable to our case.

### Testing the relative role of oceanographic connectivity in explaining genetic differentiation

Using maximum-likelihood population-effects (MLPE) linear mixed models, we tested the prediction of genetic differentiation by multigeneration connectivity and centrality separately for each species, study, genetic marker, and differentiation index. This dataset-to-dataset approach is essential as genetic differentiation data cannot be directly compared (Gouvêa et al., 2023, Legrand et al., 2022). Furthermore, the utilisation of MLPE models accounted for the non-independence of pairwise comparisons, a distinctive characteristic of genetic differentiation (Jahnke et al., 2022, Legrand et al., 2022, Selkoe et al., 2016). We constructed three independent MLPE models per dataset: 1) a Spatial Model (SM) considering a random effect at the sampling site level, 2) a Connectivity Model (CM) incorporating the distance-transformed symmetric probability of multigeneration connection as a fixed effect and sampling locality level as a random effect, and 3) a Connectivity and Centrality Model (CCM) incorporating pairwise difference of harmonic centrality as fixed effects over the previous CM. We assessed the performance of the models by comparing predicted and observed genetic differentiation with a linear model, from which AIC, goodness of fit (adjusted R^2^), and p-values were extracted. Datasets with fewer than eight pairs of sampling sites were excluded (i.e., 17 datasets, Gouvêa et al., 2022, Jenkins & Quintana-Ascencio, 2020). To account for null probabilities of connection in log-transformations, we added the minimum numerical value of R computing language (e.g., 9e-324). This adjustment effectively generated infinite distances numerically for unreachable pairs of sites.

All analyses were performed in R (R Development Core Team, 2018).

## Results

Our systematic review identified 49 studies that provided pairwise population genetic differentiation data. This resulted in 84 datasets, each representing a unique combination of a study, a species, a genetic marker, and a differentiation index (see SI Table S2 for further information). In total, it encompasses 34 species, 5 markers (isozyme, microsatellites, chloroplast DNA, mitochondrial DNA, and nuclear DNA) and 7 differentiation indices (Dst, Fst, Gst, JostD, Nei, Nst, and φst).

The dataset consists of a total of 662 populations sampled across the globe (Figure 1), with microsatellites (55.9 %) and Fst (69.0 %) being the most commonly used. On average, each dataset includes 16.7 ± 2.5 populations, with an average distance of 843.3 ± 191.1 km between them. Most of the sampling (77.4 %) was conducted within a single marine realm, primarily in the Temperate Northern Atlantic (TNA, 45.2 %), the Temperate Northern Pacific (TNP, 32.1 %), and the Temperate Australasia (TA, 16.7 %) (Figure 1, Spalding et al., 2007). The dataset exhibits significant variability in terms of spatial sampling coverage. For instance, 42 populations of *Macrocystis pyrifera* were sampled across 1,2934 km and 41° of latitude extent (Assis et al., 2023). In contrast, 21 populations of *Laminaria digitata* were sampled within 210 km, encompassing a latitudinal extent of 1.4° (Robuchon et al., 2014).

A significant linear fit between observed and predicted genetic differentiation was found for 82.1% of the Spatial Model (SM), 89.6 % of the Connectivity Model (CM) and 92.6 % of the Connectivity and Centrality Model (CCM). The average R^2^ of SMs was 0.30 ± 0.06, which was improved to 0.47 ± 0.06 with the CM and to 0.49 ± 0.07 with the CCM. The CCM displayed the lowest Akaike Information Criterion (AIC) value for 76.1% of the datasets, minimising information loss with a mean relative likelihood of 0.86 ± 0.07 (compared to 0.22 ± 0.09 for the SM and 0.36 ± 0.09 for the CM, Figure 2 and SI Table S3). The CCM stands as the optimal predictor of genetic differentiation in marine forests. The significant R^2^ values of the CCMs ranged between 0.06 (ID 45, *Sargassum polycystum*, Chan et al., 2013) and 0.95 (ID 8, *Ecklonia radiata*, Coleman et al., 2020). Only the number of sampling sites significantly affected CCM goodness of fit (linear model fits with R^2^ = 0.09, p-value = 0.01), and not the species or study characteristics (SI Table S5). The R^2^ improvement of CCM over SM spanned from -0.14 (ID 5, *Ishige okamurae*, Lee et al., 2012) to 0.92 (ID 29, *Laminaria Hyperbora*, Robuchon et al., 2014). In 88.1 % of the tests, the use of connectivity and centrality information yielded a noticeable gain in R^2^, compared to the spatial sampling effect (as depicted on Figure 3 with *Cystoseira tamariscifolia*; Bermejo et al., 2018). On average, it provided an additional explanation of 19.6 ± 5.6 % of the genetic variance, and this value reached 53.6 % for the top 10% of the dataset (Q0.9 = 0.54 ΔR^2^, Figure 2, SI Table S3).

**Figure 2:**
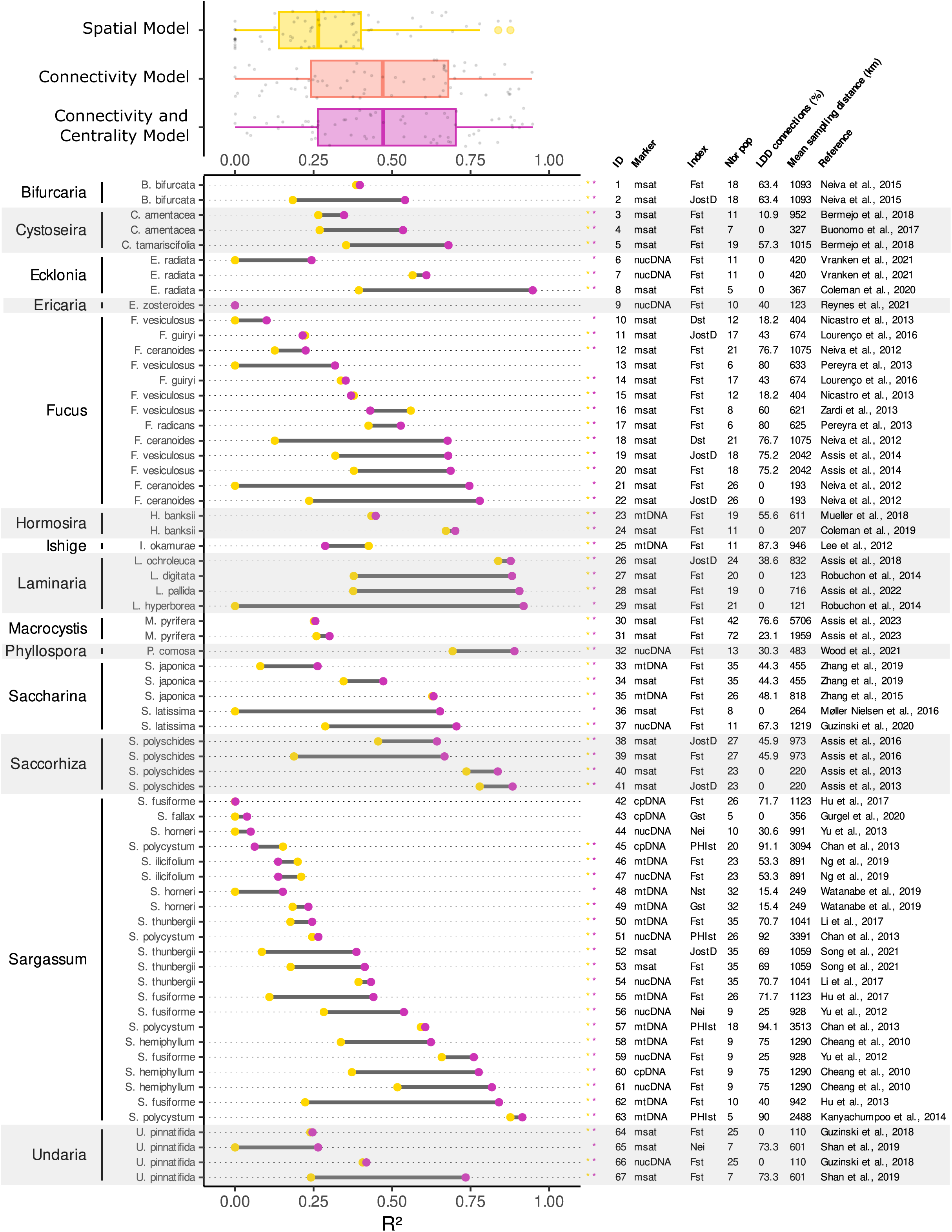
The relative role of oceanographic connectivity in explaining genetic differentiation. Boxplots illustrate the distribution of goodness of fit (R^2^) between observed genetic differentiation and predicted genetic differentiation using the Spatial Model (SM), the Connectivity Model (CM), and the Connectivity and Centrality Model (CCM). Dataset-specific R^2^ is highlighted for SM (yellow dots) and CCM (purple dots), while the difference of R^2^ between these two models (ΔR^2^) is denoted with a grey bar. Significant models (p-value < 0.05) are indicated with colored asterisks (i.e., yellow for SM, purple for CCM). Information about the modelled species and its genera, the genetic marker (Marker), the genetic differentiation index (Index), the number of sampled populations (Nbr pop, at the 8.5 km spatial resolution of the biophysical model), the percentage of connection between sampling site promoted by LDD (LDD connections), the mean distance between sampled populations (Mean sampling distance) and the reference is notified for each dataset. Genetic markers are shown as msat (microsatellites), cpDNA (chloroplast DNA, i.e., rbcL, plastidal RuBisCO spacer), mtDNA (mitochondrial DNA, i.e., CO1, COX1, COX3, Trn spacers), and nucDNA (nuclear DNA, i.e., SNPS, ISSR primers, SRAP primers, ITS2). ID permits to refer to a given dataset in the main text. On each boxplot, the central mark, the bottom and top edges indicate the median, the 25th and the 75th percentiles, respectively. The whiskers extend to the most extreme data points not considered outliers, and the outliers are plotted individually using a dot symbol.

**Figure 3:**
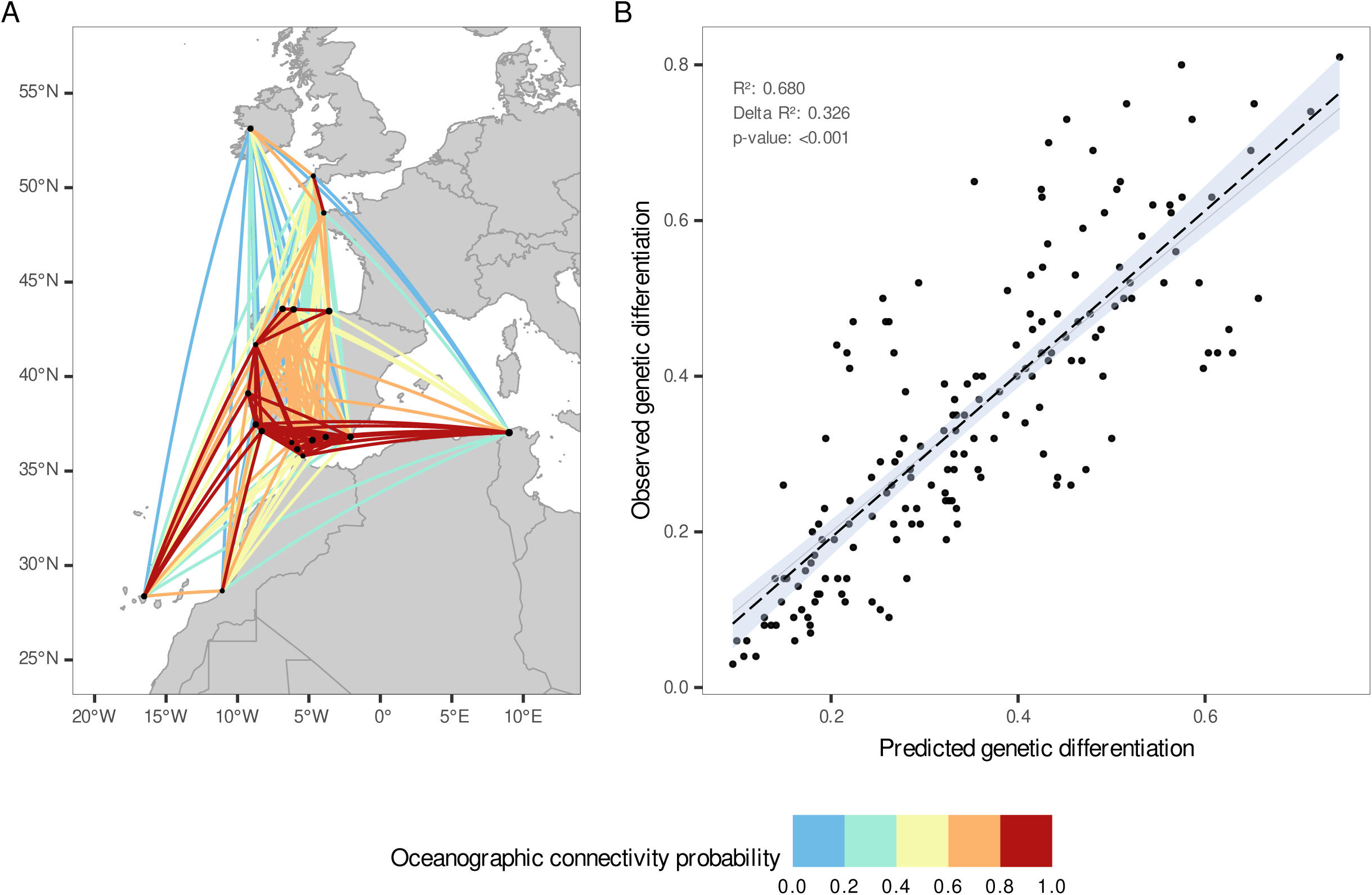
Connectivity networks and fit between predicted and observed genetic differentiation for *Cystoseira tamariscifolia* (ID 5, Bermejo et al., 2018). Panel A depicts the network’ relative probability of oceanographic connectivity between sampled populations. Note that multi-generations connections are depicted, and not stepping-stone pathways. Panel B displays the predicted genetic differentiation with CCM against the observed genetic differentiation. Dotted black lines indicate the linear regression trend and blue shading represents the corresponding 95 % confidence interval.

Considering LDD, contrary to a simplistic approach of a fixed PD value in the CCMs (Figure 4a, SI Table S4), allowed an additional goodness of fit of 0.04 ± 0.04, improving R^2^ in 56.7 % of the datasets and reducing AIC in 56.7 % of the datasets, with a mean relative likelihood of 0.66 ± 0.11. LDD events not only cross-continent connections (e.g., South America to Africa and Oceania, North America to Europe) or connections between isolated islands and continents (e.g., Azores, Iceland, Guam, Okinawa; Figure 4c), but also increased the number of connected populations by more than half (i.e., from 47% for fixed PD to 99% for LDD). Additionally, populations required an average of 6.15 ± 0.18 steps to be connected with LDD, compared to 25.1 ± 0.83 steps without it (Figure 4b). Consequently, LDD allows populations to connect, on average, 7.22 ± 0.26 times faster (SI Table S3, S4).

**Figure 4:**
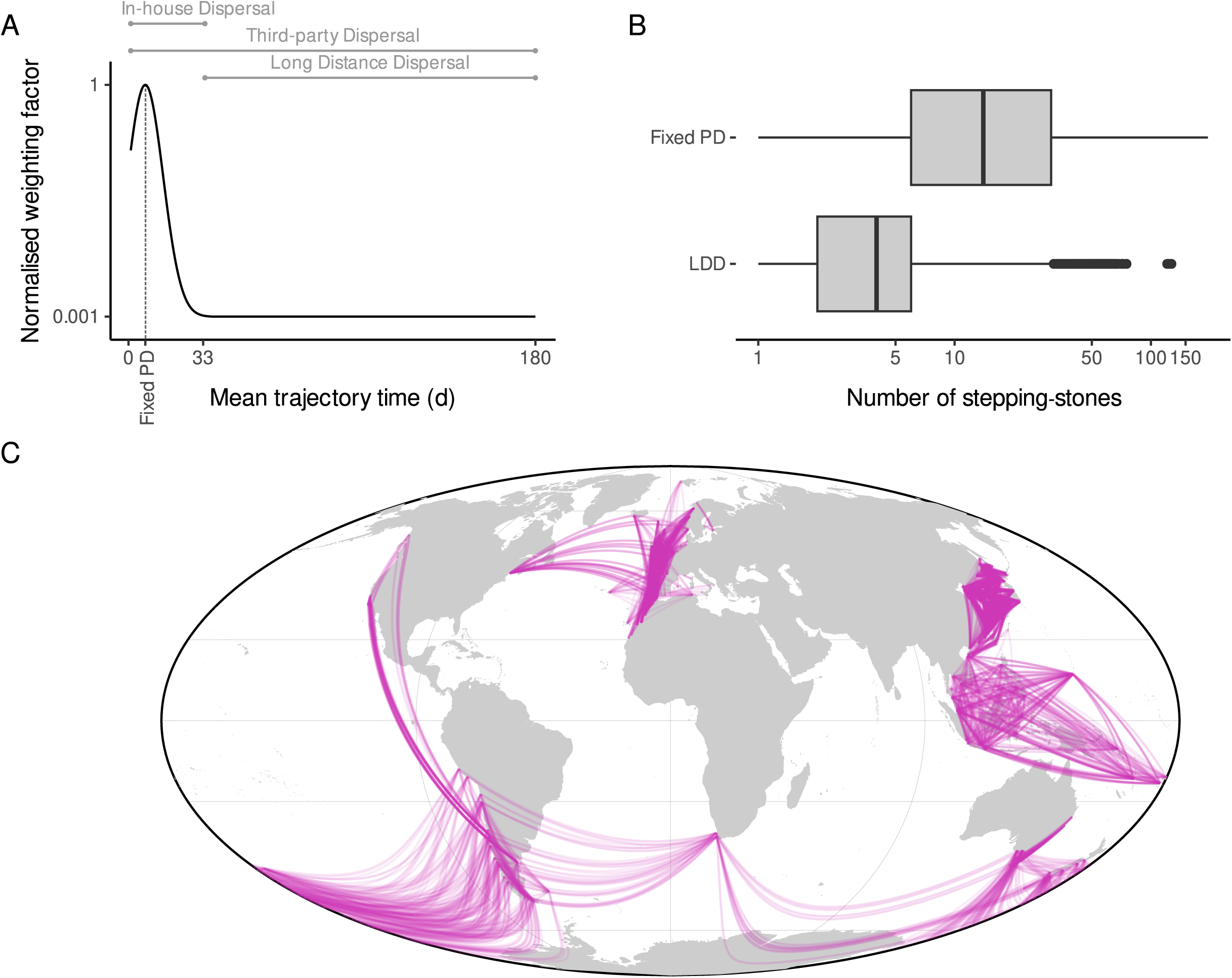
Marine forest dispersal capacity and LDD consideration. Panel A displays the weighting factor of probabilities of connections regarding the mean trajectory time between a pair of sites, given by a normal distribution fitted on in-house dispersal capacity (μ = mean of dispersal range time, 7.43 d, see SI Table S1). LDD events start at 33 days (i.e., maximal in-house dispersal capacity, Deysher et al., 1981) with a unique weighting factor of 0.01, which corresponds to the proportion of LDD events observed using population assignment tests (D’Aloia et al., 2022). Panel B indicates the distribution of the number of stepping-stones between sampling sites, considering the fixed PD case (i.e., 7.43 d) and the full dispersal capacity considering LDD. The whiskers extend to the most extreme data points not considered outliers. Panel C shows the connections between sampling sites permitted by LDD. It represents 52.6 % of the total number of connections. Note that multi-generations connections are depicted and not stepping-stone pathways. On each boxplot, the central mark, the bottom and the top edges indicate the median, the 25th and the 75th. The whiskers extend to the most extreme data points not considered outliers, and the outliers are plotted individually using a dot symbol.

## Discussion

Our results represent the first attempt to unravel the relative contribution of oceanographic connectivity and centrality on the global intra-specific diversity patterns of key ecosystem structuring species of marine forests. We show that independently of the species, genetic marker, or differentiation index, our models explained a large part (49.5 %) of the genetic differentiation variance. Specifically, in 92.6 % of the datasets, spatial dependent processes explained 29.9 % of genetic differentiation and oceanographic connectivity coupled with centrality explained an additional 19.6 %. The consideration of potential LDD permitted capturing cross-continent connections, increasing the number of connected populations from 46.1 % with fixed PD values to 99.3 %. Neglecting LDD, as commonly used (e.g., Assis et al., 2018, 2022, Buonomo et al., 2017, Pereyra et al., 2013, Reynes et al., 2021), led to higher stepping-stones required to connect populations and a higher proportion of unconnected populations, a pattern that is not supported by independent genetic differentiation estimates. These results have significant implications for marine forests’ biogeography and evolution and further highlight the importance of considering oceanographic connectivity and centrality in conservation, management, and climate change research.

The spatial models considering the effect of spatial sampling alone were significant for 82.1 % of the datasets, explaining 29.9 % of the overall variance in genetic differentiation. These models reflect various spatially dependent processes underlying the distribution of marine forest species. Specifically, habitat discontinuities caused by variations in substrate types or habitat fragmentation (Robuchon et al., 2014) and physical barriers, such as isolated seascape features like islands, seamounts, or peninsulas. Such discontinuities and barriers can lead to intrinsic reproductive isolation, with offspring exhibiting reduced viability, fitness, survival rates, and reproductive success, or even sterility, regardless of the connectivity levels between populations (Bierne et al., 2011, Guillemin et al., 2016). Reproductive isolation can also occur between populations that belong to different ecotypes (Coleman et al., 2019, Zardi et al., 2013). Selective pressures like temperature (Neiva et al., 2015, Nicastro et al., 2013, Vranken et al., 2021, Wood et al., 2021, Zhang et al., 2019), salinity (Møller Nielsen et al., 2016), light intensity (Reynes et al., 2021) and immersion (Robuchon et al., 2014) can lead to local adaptation (including differences in reproduction timing) and thus genetic differentiation levels among the sampled populations. Accordingly, extreme environmental events, such as marine heat waves, can potentially reduce effective population sizes, produce genetic bottlenecks due to drift, and alter local gene pools (Coleman et al., 2020, Gurgel et al., 2020). Drift also promotes gene surfing, a process typically occurring during post-glacial range shifts, in which low-frequency alleles rapidly spread along the leading edge, shaping spatial patterns of genetic diversity (Assis et al., 2016, 2018, Bermejo et al., 2018, Song et al., 2021). While the spatial model implicitly considers the aforementioned processes, explaining more than one-third of the genetic differentiation, their relative influence remains unclear.

The models considering oceanographic connectivity and centrality estimates were significant in 92.6% of the cases and explained an additional 19.6 % of genetic differentiation data, which builds upon previous research performed on mangrove forests (Gouvea et al., 2023), and specific geographical areas, such as the Mediterranean Sea (Legrand et al., 2022). This improvement in explaining genetic differentiation likely resulted from the oceanographic data used in biophysical modelling that was able to resolve key meso-scale processes (e.g., fronts, bifurcations and eddies) occurring within the range of marine forests’ distribution. Such processes, driven by the long-term patterns of ocean currents, might have determined the extent to which sampled populations are connected, by promoting unidirectional or bidirectional gene flow or by structuring oceanographic barriers that might have isolated populations and contributed to increased genetic differentiation (Liang et al., 2022). Accordingly, considering multigenerational stepping-stone connectivity and the centrality degree of populations was particularly important in explaining genetic differentiation. On the one hand, stepping-stone connectivity accounted for gene flow events occurring across multiple generations and along the complex pathways involved (Buonomo et al., 2017). Importantly, direct connectivity between discrete sampled populations is very unlikely (Legrand et al., 2022), especially when these are distributed across large spatial scales, such as in many of our datasets. On the other hand, centrality accounted for the relative importance or prominence of specific populations, which can serve as connectivity hubs and significantly facilitate gene flow. High centrality populations (i.e., hubs for genetic connectivity), more easily accessible, can serve as stepping-stones or bridges between other populations, thus reducing genetic differentiation levels. Conversely, low centrality populations are associated with regions of genetic discontinuities, limiting gene flow and contributing to increased differentiation (Gouvea et al., 2023; Rozenfeld et al., 2008). Mapping the relative centrality of marine forest populations can inform conservation and management strategies in the face of changing environmental conditions and ongoing anthropogenic impacts (Gouvea et al., 2023). In particular, the conservation of hubs for connectivity with higher centrality, facilitating LDD events or bridging populations across large water masses, can help safeguard the genetic integrity and resilience of populations (Abecasis et al., 2023; Assis et al., 2021).

In our study, we conducted a comprehensive assessment of marine forests’ population connectivity. Instead of setting a fixed or multiple pelagic duration values corresponding to marine forest species’ dispersal ability, our approach aimed at capturing both the intra-specific variability in dispersal capacity and the LDD facilitated by the detachment of fragments or thalli. We consider the occurrence of LDD to have a probability of 0.1 % (d’Aloia et al., 2022), in addition to the already minimal probability of propagules connecting distant sites, as observed by the skewed dispersal kernel of mangroves described at global scale (Gouvêa et al., 2023). LDD events, which may span thousands of km (Gouvêa et al., 2023, Smith et al., 2018), played a crucial role in reducing the number of stepping-stone iterations. On average, it led to a seven-fold decrease in the number of steps required to connect populations, driving to an average shortest path length of ∼ 6 steps, which is lower than previous estimations performed at local scales (a few dozen of steps in Buonomo et al., 2017, Boulanger et al., 2020, Legrand et al., 202). Despite low probability, LDD not only facilitated the connection of distant and isolated populations but also promoted links for over half of the observed pairs of populations, highlighting its disproportionate influence on contemporary gene flow (Jordano et al., 2017, Nathan et al., 2006). Failing to consider LDD and, therefore, all putative stepping-populations, would overlook critical connections that support the relatively low genetic differentiation of sampled populations (e.g., average Fst: 0.34 ± 0.01), even when vast oceanic distances separate them (Tavares et al., 2023). However, it is hard to adequately capture the intensity at which such sporadic events shape and disrupt the local structure of gene pools. We have forced a 0.1% threshold to penalise the estimated probabilities of connectivity beyond the maximum reported propagule durations, while others have reported even more restrictive ratios (Smith et al., 2018). Despite the uncertainty, models incorporating LDD showed an average model improvement of 3.8 % of variance over those simplistically considering a unique infinite distance.

The connectivity and centrality models, attributed to spatial dependent processes, explained 49.5 % of the variance in genetic differentiation, of which 19.6 % is explicitly influenced by contemporary connectivity. The remaining unexplained variance may be attributed to additional mechanisms not considered, mainly at the population dynamic level. The Allee effect, for instance, predicts higher fitness and survival of individuals in larger populations (Stephens et al., 1999). Accordingly, in small populations, the reproductive potential may be disrupted as fewer propagules are released, thus hindering dispersal and settlement capacity. From a network perspective, such populations act as sinks and not as sources of propagules. Conversely, when large populations reach their carrying capacity, settlement or mating processes for immigrants can be prevented by density barriers (Del Monte-Luna et al., 2004, Water et al., 2013). This means dispersal can be effective, but habitats are limited or unavailable for incoming individuals (from any form, spore, gamete, or detached fragment). As such, well-established populations act as sources and not as sinks of propagules. Moreover, marine forest population sizes can vary over time due to the balance between mortality (which can be induced by anthropogenic impacts), self-recruitment and immigration (caused directly by connectivity for sessile organisms). Such meta-population dynamics, not represented in our modelling approach, can alter or reinforce realised genetic connectivity at each successive generation, playing an important role not only in contemporary genetic differentiation patterns but also in eco-evolutionary processes such as postglacial recolonisation and range expansion.

The unexplained variance in genetic differentiation could also be induced by uncertainties and limitations of the biophysical model. As it permits us to evaluate connectivity patterns at global scale, the resolution of the velocity fields used in our model (1/16°) can leave nearshore dynamics unresolved. Such submesoscale physical processes (McWilliams et al., 2016) have a crucial role in the trajectories of propagule dispersing for a few days (typically in-house dispersal, i.e., gamete, zygote and/or spore in macroalgae), possibly affecting local connectivity patterns (Saint-Amand et al., 2023, Ward et al., 2023). Similarly, uncertainty also remains in third-party dispersal, as we have not considered the effect of wind and waves on drift velocities of macroalgae rafts when simulating LDD (Podlejski et al., 2023). Dispersal capacity reported for marine forests is highly variable in the literature, with a lack of species-specific information (i.e., we have insufficient material for 79 % of the species compiled in our dataset). The normal distribution aiming to represent in-house dispersal according to observed dispersal capacity, therefore, results in a generalized function whose precision may vary for the species under consideration. This calls into question the reliability of dispersal capacity as a parameter of the biophysical model and suggests that a better understanding of this biological trait could contribute to a more comprehensive modelling of dispersal (Jahnke et al., 2022). When establishing links between populations, searching for the shortest path does not accumulate all the possible paths nor disentangle implicit connections between populations sharing a common ancestor (McRae & Beier, 2007, Ser-Giacomi et al., 2021). These features are essential to evaluate coalescent connectivity, which has been shown to better predict gene flow (Legrand et al., 2022). Such connections could be implemented in future studies. Ultimately, focusing on species-specific distribution through a deeper consideration of species dispersal traits and habitats (Assis et al., 2018, 2022, Mari et al., 2022) could bring an improvement for this approach.

Recent calls for global biodiversity conservation have emphasised the need to consider the preservation of the genetic diversity of wild populations (Laikre et al., 2020), yet, conservation and management actions have largely overlooked it (Andrello et al., 2022; Carvalho et al., 2017). Our results show the influence of spatial-dependent processes and oceanographic connectivity on the contemporary intra-specific diversity of marine forests. This means that there are areas with specific spatial features worthy of conservation (e.g., as gene conservation units; Andrello et al., 2022). Accordingly, mapping centrality and isolation for the designation of Marine Protected Areas can find bridging marine forests’ populations that facilitate gene flow or isolated populations that could shelter unique genetic assemblage (Assis et al., 2018, 2021; Abecasis et al., 2023). Such an approach could guide the implementation of the post-2020 framework, which considers the conservation of 30% of the oceans through well-connected MPAs (Dinerstein et al., 2019; Magris et al., 2018). Beyond MPA designation, oceanographic connectivity patterns can also be utilised in marine forest aquaculture spatial planning (Brackel et al., 2021). This would facilitate the design of management strategies that minimise the risk of genetic introgression from cultivated populations into natural populations, thereby preserving the unique genetic characteristics and adaptations of each population. In conclusion, our findings underscore the significance of considering oceanographic connectivity and centrality in safeguarding the genetic integrity and resilience of ecosystem structuring species of brown macroalgae.

## Acknowledgements

This work was funded by Portuguese National Funds from FCT - Foundation for Science and Technology - through UIDB/04326/2020, UIDP/04326/2020, LA/P/0101/2020, PTDC/BIA-CBI/6515/2020, and the Individual Call to Scientific Employment Stimulus 2022.00861.CEECIND.

