## Supplementary Table S1, S2, S3, S4 and S5 for "Unravelling the role of oceanographic connectivity in intra-specific diversity of marine forests at global scale"

This file includes:

- Tables S1-S5
- SI references

Table S1: Dataset of macro-algae dispersal capacity, including Ochrophyta, Rhodophyta and Chlorophyta species.

| Species characteristics | | | | Dispersal duration estimation | | Study characteristics | | |
|---|---|---|---|---|---|---|---|---|
| **Species** | **Dispersal medium** | **Dispersal feature** | **Dispersal category** | **Minimum (d)** | **Maximum (d)** | **Method** | **Method feature** | **Reference** |
| *Sargassum Spp* | propagules | spore dispersal | in-house | 0 | 1 | field experiment | capture of propagule along seabed | Kendrick et al., 1995 |
| *Hormosira banksii* | fragment | fragment viability | third-party | 7 | 60 | lab experiment | reproductive viability of fragment | McKenzie et al., 2008 |
| *Durvillaea antarctica* | thalli | rafting | third-party | 20 | 65 | field estimation | growth of epifauna (*Lepas australis*) | Frazer et al., 2011 |
| *Yuzurua poiteaui* | fragment | attachment ability | third-party | 0 | 2 | lab experiment, field experiment | time needed for a fragment to attach to substrate | Heren et al., 2013 |
| *Gracilaria salicornia* | fragment | fragment longevity | third-party | 20 | 60 | lab experiment | regenerative capacity of fragment | Bierwagen et al., 2015 |
| *Ulva intestinalis* | spore | spore dispersal | in-house | 1 | 5 | lab experiment | spore time of survival | Hoffmann et al., 1989 |
| *Gymnogongrus durvillei* | spore | spore dispersal | in-house | 1 | 6 | lab experiment | spore time of survival | Hoffmann et al., 1989 |
| *Gymnogongrus furcellatus* | spore | spore dispersal | in-house | 1 | 5 | lab experiment | spore time of survival | Hoffmann et al., 1989 |
| *Nothogenia fastigiata* | spore | spore dispersal | in-house | 1 | 5 | lab experiment | spore time of survival | Hoffmann et al., 1989 |
| *Ulva rigida* | spore | spore dispersal | in-house | 1 | 7 | lab experiment | spore time of survival | Hoffmann et al., 1989 |
| *Gelidium chilense* | spore | spore dispersal | in-house | 1 | 11 | lab experiment | spore time of survival | Hoffmann et al., 1989 |
| *Pyropia columbina* | spore | spore dispersal | in-house | 1 | 11 | lab experiment | spore time of survival | Hoffmann et al., 1989 |
| *Lessonia nigrescens* | spore | spore dispersal | in-house | 1 | 5 | lab experiment | spore time of survival | Hoffmann et al., 1989 |
| *Pterygophora californica* | spore | spore dispersal | in-house | 0 | 5 | lab experiment | spore swimming | Reed et al., 1992 |
| *Macrocystis pyrifera* | spore | spore dispersal | in-house | 0 | 4 | lab experiment | spore swimming | Reed et al., 1992 |
| *Laminaria hyperborea* | spore | spore dispersal | in-house | 0 | 6 | lab experiment | spore time before settle | Kain et al., 1964 |
| *Sargassum muticum* | germlings | attachment ability | in-house | 2 | 33 | lab experiment | attachment ability after days in suspension | Deysher et al., 1981 |
| *Macrocystis pyrifera* | thalli | rafting | third-party | 16 | 76 | field experiment | change in length of blades following detachment | Hobday et al., 2000 |
| *Macrocystis pyrifera* | spore | spore dispersal | in-house | 0 | 4 | lab experiment | spore settlement rate | Gaylord et al., 2002 |
| *Undaria pinnatifida* | spore | spore dispersal | in-house | 0 | 14 | lab experiment | spore viability | Forrest et al., 2000 |
| *Fucus distichus* | zygote | zygote dispersal | in-house | 0 | 2 | lab experiment | zygote attachment | Coleman et al., 2005 |
| *Macrocystis pyrifera* | thalli | rafting | third-party | 0 | 260 | field observation | estimation of dispersal duration | Smith et al., 2002 |
| *Durvillaea antarctica* | thalli | rafting | third-party | 0 | 260 | field observation | estimation of dispersal duration | Smith et al., 2002 |
| *Sargassum ringgoldianum* | thalli | rafting | third-party | 1 | 167 | field experiment | mark and recapture | Ida et al., 1967 |

| Species characteristics | | | | Dispersal duration estimation | | Study characteristics | | |
|---|---|---|---|---|---|---|---|---|
| **Species** | **Dispersal medium** | **Dispersal feature** | **Dispersal category** | **Minimum (d)** | **Maximum (d)** | **Method** | **Method feature** | **Reference** |
| *Ecklonia maxima* | thalli | rafting | third-party | 0 | 150 | field observation | estimation of dispersal duration | Arnaud et al., 1976 |
| *Ascophyllum nodosum* | thalli | rafting | third-party | 0 | 43 | field observation | algea raft survival | Ingólfsson et al., 1998 |

Table S2: Sampling characteristics of datasets. Species, genetic marker (Marker), genetic differentiation index (Diff.Index), dataset speciality (Spec), number of populations (n), the maximum distance between sampling sites in km (Samp.Ext), the mean distance between sampling sites in km (Samp.Mean.Dist), the maximum latitude extent between sampling sites in degree (Lat.Ext), the sampled realm (Realm) and the dataset reference are reported. Genetic markers are shown as isozyme, msat (microsatellites), cpDNA (chloroplast DNA, i.e. rbcL, plastidal RuBisCO spacer), mtDNA (mitochondrial DNA, i.e. CO1, COX1, COX3, Trn spacers), and nucDNA (nuclear DNA, i.e. SNPS, ISSR primers, SRAP primers, ITS2). Realms (Spalding et al., 2007) are abbreviated as TNA (Temperate Northern Atlantic), TA (Temperate Australasia), TNP (Temperate Northern Pacific), CIP (Central Indo-Pacific), A (Arctic), WIP (Western Indo-Pacific), EIP (Eastern Indo-Pacific), TSAf (Temperate Southern Africa), TSAm (Temperate South America), SO (Southern Ocean).

| Species | Marker | Diff.Index | Spec | n | Samp.Ext | Samp.Mean.Dist | Lat.Ext | Realm | Reference |
|---|---|---|---|---|---|---|---|---|---|
| Bifurcaria bifurcata | msat | Fst | | 18 | 2829.12 | 1091.99 | 25.3 | TNA | Neiva et al., 2015 |
| Bifurcaria bifurcata | msat | JostD | | 18 | 2829.12 | 1091.99 | 25.3 | TNA | Neiva et al., 2015 |
| Cystoseira amentacea | msat | Fst | | 11 | 1893.49 | 953.36 | 6.5 | TNA | Bermejo et al., 2018 |
| Cystoseira amentacea | msat | Fst | | 7 | 572.89 | 327.67 | 3.5 | TNA | Buonomo et al., 2017 |
| Cystoseira mediterranea | msat | Fst | | 4 | 322.11 | 209.64 | 2.9 | TNA | Bermejo et al., 2018 |
| Cystoseira tamariscifolia | msat | Fst | | 19 | 2818.33 | 1014.33 | 24.8 | TNA | Bermejo et al., 2018 |
| Ecklonia radiata | nucDNA | Fst | adaptive | 11 | 827.28 | 419.35 | 6.5 | TA | Vranken et al., 2021 |
| Ecklonia radiata | nucDNA | Fst | neutral | 11 | 827.28 | 419.35 | 6.5 | TA | Vranken et al., 2021 |
| Ecklonia radiata | msat | Fst | beforevsafter | 5 | 726.74 | 366.3 | 6.5 | TA | Coleman et al., 2020 |
| Ecklonia radiata | msat | Fst | before | 4 | 726.74 | 399.91 | 6.5 | TA | Coleman et al., 2020 |
| Ecklonia radiata | msat | Fst | after | 4 | 605.23 | 337.92 | 5.4 | TA | Coleman et al., 2020 |
| Ericaria zosteroides | nucDNA | Fst | | 10 | 282.54 | 122.89 | 0.9 | TNA | Reynes et al., 2021 |
| Fucus ceranoides | msat | Fst | | 26 | 634.64 | 193.8 | 2 | TNA | Neiva et al., 2012 |
| Fucus ceranoides | msat | JostD | | 26 | 634.64 | 193.8 | 2 | TNA | Neiva et al., 2012 |
| Fucus ceranoides | msat | Fst | | 21 | 3216.19 | 1076.65 | 25.7 | TNA | Neiva et al., 2012 |
| Fucus ceranoides | msat | Dst | | 21 | 3216.19 | 1076.65 | 25.7 | TNA | Neiva et al., 2012 |
| Fucus distichus | msat | Fst | | 23 | 125.29 | 125.29 | 0.5 | TNA | Coleman et al., 2005 |
| Fucus guiryi | msat | Fst | | 17 | 1815.86 | 673.18 | 15.4 | TNA | Lourenço et al., 2016 |
| Fucus guiryi | msat | JostD | | 17 | 1815.86 | 673.18 | 15.4 | TNA | Lourenço et al., 2016 |
| Fucus radicans | msat | Fst | | 6 | 1239.85 | 626.18 | 10.9 | TNA | Pereyra et al., 2013 |
| Fucus spiralis | msat | Fst | | 8 | 128.97 | 128.97 | 0.5 | TNA | Coleman et al., 2005 |
| Fucus vesiculosus | msat | Fst | | 18 | 5438.93 | 2045.38 | 29 | TNA | Assis et al., 2014 |
| Fucus vesiculosus | msat | JostD | | 18 | 5438.93 | 2045.38 | 29 | TNA | Assis et al., 2014 |
| Fucus vesiculosus | msat | Fst | | 6 | 1264.14 | 634.77 | 11.1 | TNA | Pereyra et al., 2013 |
| Fucus vesiculosus | msat | Fst | | 12 | 926.38 | 403.43 | 8.3 | TNA | Nicastro et al., 2013 |
| Fucus vesiculosus | msat | Dst | | 12 | 926.38 | 403.43 | 8.3 | TNA | Nicastro et al., 2013 |
| Fucus vesiculosus | msat | Fst | | 8 | 1051.75 | 621.52 | 8.7 | TNA | Zardi et al., 2013 |
| Halidrys dioica | isozyme | Nei | | 5 | 81.93 | 81.93 | 0.6 | TNP | Lu et al., 1994 |

| Species | Marker | Diff.Index | Spec | n | Samp.Ext | Samp.Mean.Dist | Lat.Ext | Realm | Reference |
|---|---|---|---|---|---|---|---|---|---|
| Hormosira banksii | msat | Fst | | 11 | 437.29 | 207.12 | 3.6 | TA | Coleman et al., 2019 |
| Hormosira banksii | mtDNA | Fst | | 19 | 1831.75 | 610.1 | 15.3 | TA | Mueller et al., 2018 |
| Ishige okamurae | mtDNA | Fst | | 11 | 2895.97 | 946.57 | 13.3 | TNP,CIP | Lee et al., 2012 |
| Laminaria digitata | msat | Fst | | 16 | 15.39 | 15.39 | 0.1 | TNA | Brennan et al., 2014 |
| Laminaria digitata | msat | Fst | | 20 | 210.96 | 123.59 | 1.4 | TNA | Robuchon et al., 2014 |
| Laminaria hyperborea | msat | Fst | | 21 | 210.96 | 121.01 | 1.4 | TNA | Robuchon et al., 2014 |
| Laminaria ochroleuca | msat | JostD | | 24 | 2194.35 | 831.55 | 18.8 | TNA | Assis et al., 2018 |
| Laminaria pallida | msat | Fst | | 19 | 1775.57 | 714.32 | 15.1 | TSA | Assis et al., 2022 |
| Macrocystis pyrifera | msat | Fst | Southern hemisphere | 42 | 12951.53 | 5716.91 | 41 | TSA,SO,TA | Assis et al., 2023 |
| Macrocystis pyrifera | msat | Fst | Northern hemisphere | 72 | 12782.89 | 1955.76 | 99.1 | TSA,TNP | Assis et al., 2023 |
| Nereia lophocladia | nucDNA | Fst | | 7 | 67.36 | 36.78 | 0.6 | TA | Mamo et al., 2021 |
| Phyllospora comosa | nucDNA | Fst | | 13 | 1431.17 | 482.22 | 12 | TA | Wood et al., 2021 |
| Pylaiella littoralis | msat | Fst | | 6 | 166.57 | 119.58 | 0.9 | TNA | Geoffroy et al., 2015 |
| Saccharina japonica | msat | Fst | | 35 | 1613.52 | 455.48 | 11.1 | TNP | Zhang et al., 2019 |
| Saccharina japonica | mtDNA | Fst | | 35 | 1613.52 | 455.48 | 11.1 | TNP | Zhang et al., 2019 |
| Saccharina japonica | mtDNA | Fst | | 26 | 2155.96 | 819.01 | 11.7 | TNP | Zhang et al., 2015 |
| Saccharina japonica | nucDNA | Fst | | 5 | 1186.94 | 1186.94 | 10.6 | TNP | Zhao et al., 2013 |
| Saccharina japonica | nucDNA | Nei | | 5 | 1186.94 | 1186.94 | 10.6 | TNP | Zhao et al., 2013 |
| Saccharina latissima | msat | Fst | | 8 | 677.2 | 264.22 | 5.2 | TNA | Møller Nielsen et al., 2016 |
| Saccharina latissima | msat | Fst | | 5 | 143.12 | 83.97 | 0.5 | TNA | Breton et al., 2018 |
| Saccharina latissima | nucDNA | Fst | | 11 | 4270.39 | 1220.95 | 37.7 | TNA,A | Guzinski et al., 2020 |
| Saccorhiza polyschides | msat | Fst | | 27 | 3395.93 | 973.05 | 28.9 | TNA | Assis et al., 2016 |
| Saccorhiza polyschides | msat | JostD | | 27 | 3395.93 | 973.05 | 28.9 | TNA | Assis et al., 2016 |
| Saccorhiza polyschides | msat | Fst | | 23 | 517.39 | 219.53 | 4.7 | TNA | Assis et al., 2013 |
| Saccorhiza polyschides | msat | JostD | | 23 | 517.39 | 219.53 | 4.7 | TNA | Assis et al., 2013 |
| Sargassum fallax | mtDNA | Gst | before | 4 | 726.61 | 486.87 | 6.5 | TA | Gurgel et al., 2020 |
| Sargassum fallax | mtDNA | Gst | after | 5 | 552.18 | 374.59 | 5 | TA | Gurgel et al., 2020 |
| Sargassum fallax | cpDNA | Gst | before | 4 | 726.61 | 399.97 | 6.5 | TA | Gurgel et al., 2020 |
| Sargassum fallax | cpDNA | Gst | after | 5 | 726.61 | 355.51 | 6.5 | TA | Gurgel et al., 2020 |
| Sargassum fusiforme | mtDNA | Fst | | 26 | 3521.7 | 1123.17 | 19.4 | TNP,CIP | Hu et al., 2017 |
| Sargassum fusiforme | cpDNA | Fst | | 26 | 3521.7 | 1123.17 | 19.4 | TNP,CIP | Hu et al., 2017 |
| Sargassum fusiforme | mtDNA | Fst | | 10 | 2347.8 | 940.36 | 18.2 | TNP,CIP | Hu et al., 2013 |
| Sargassum fusiforme | nucDNA | Fst | | 9 | 2347.8 | 926.75 | 18.2 | TNP,CIP | Yu et al., 2012 |
| Sargassum fusiforme | nucDNA | Nei | | 9 | 2347.8 | 926.75 | 18.2 | TNP,CIP | Yu et al., 2012 |
| Sargassum hemiphyllum | nucDNA | Fst | | 9 | 2814.71 | 1290.05 | 12.8 | TNP,CIP | Cheang et al., 2010 |
| Sargassum hemiphyllum | cpDNA | Fst | | 9 | 2814.71 | 1290.05 | 12.8 | TNP,CIP | Cheang et al., 2010 |
| Sargassum hemiphyllum | mtDNA | Fst | | 9 | 2814.71 | 1290.05 | 12.8 | TNP,CIP | Cheang et al., 2010 |
| Sargassum horneri | mtDNA | Gst | | 32 | 1034.59 | 248.86 | 2.8 | TNP | Watanabe et al., 2019 |

| Species | Marker | Diff.Index | Spec | n | Samp.Ext | Samp.Mean.Dist | Lat.Ext | Realm | Reference |
|---|---|---|---|---|---|---|---|---|---|
| Sargassum horneri | mtDNA | Nst | | 32 | 1034.59 | 248.86 | 2.8 | TNP | Watanabe et al., 2019 |
| Sargassum horneri | nucDNA | Nei | | 10 | 2235.42 | 988.73 | 17.9 | TNP,CIP | Yu et al., 2013 |
| Sargassum ilicifolium | nucDNA | Fst | | 23 | 2971.53 | 890.91 | 15.2 | TNP,CIP | Ng et al., 2019 |
| Sargassum ilicifolium | mtDNA | Fst | | 23 | 2971.53 | 890.91 | 15.2 | TNP,CIP | Ng et al., 2019 |
| Sargassum polycystum | mtDNA | PHIst | | 5 | 4096.54 | 2482.5 | 35 | CIP,WIP | Kanyachumpoo et al., 2014 |
| Sargassum polycystum | nucDNA | PHIst | | 26 | 9376.05 | 3389.92 | 36.4 | CIP,WIP,EIP | Chan et al., 2013 |
| Sargassum polycystum | cpDNA | PHIst | | 20 | 9102.64 | 3092.15 | 36.4 | CIP,WIP | Chan et al., 2013 |
| Sargassum polycystum | mtDNA | PHIst | | 18 | 9137.63 | 3511.45 | 36.4 | CIP,WIP | Chan et al., 2013 |
| Sargassum thunbergii | nucDNA | Fst | | 35 | 2816.12 | 1040.8 | 18.2 | TNP | Li et al., 2017 |
| Sargassum thunbergii | mtDNA | Fst | | 35 | 2816.12 | 1040.8 | 18.2 | TNP | Li et al., 2017 |
| Sargassum thunbergii | msat | Fst | | 35 | 2808.35 | 1059.53 | 18.3 | TNP | Song et al., 2021 |
| Sargassum thunbergii | msat | JostD | | 35 | 2808.35 | 1059.53 | 18.3 | TNP | Song et al., 2021 |
| Undaria pinnatifida | msat | Fst | native | 3 | 906.92 | 617.99 | 8.1 | TNP | Shan et al., 2019 |
| Undaria pinnatifida | msat | Fst | invasive | 7 | 1135.04 | 602.64 | 6.4 | TNA | Shan et al., 2019 |
| Undaria pinnatifida | msat | Nei | native | 3 | 906.92 | 617.99 | 8.1 | TNP | Shan et al., 2019 |
| Undaria pinnatifida | msat | Nei | invasive | 7 | 1135.04 | 602.64 | 6.4 | TNA | Shan et al., 2019 |
| Undaria pinnatifida | nucDNA | Fst | | 25 | 192.07 | 109.98 | 1.2 | TNA | Guzinski et al., 2018 |
| Undaria pinnatifida | msat | Fst | | 25 | 192.07 | 109.98 | 1.2 | TNA | Guzinski et al., 2018 |

Table S3: Performance of Spatial Model (SM), Connectivity Model (CM) and Connectivity and Centrality Model (CCM), using the full consideration of dispersal abilities. Species, genetic marker, genetic differentiation index (Diff.Index), dataset speciality (Spec), number of pairs of sites (Pairs), number of pairs of sites connected (Pairs.Co) and dataset reference are reported. Performance of models are given with Akaike information criterion (AIC), significance levels (p-value) and the goodness of fit ($R^2$).

| | | | | | | **AIC** | | | ***p*-value** | | | **$R^2$** | | | |
|---|---|---|---|---|---|---|---|---|---|---|---|---|---|---|---|
| **Species** | **Marker** | **Diff.Index** | **Spec** | **Pairs** | **Pairs.Co** | **SM** | **CM** | **CCM** | **SM** | **CM** | **CCM** | **SM** | **CM** | **CCM** | **Reference** |
| Bifurcaria bifurcata | msat | Fst | | 153 | 153 | -140.85 | -141.35 | -143.76 | 0 | 0 | 0 | 0.39 | 0.39 | 0.4 | Neiva et al., 2015 |
| Bifurcaria bifurcata | msat | JostD | | 153 | 153 | -244.54 | -322.84 | -332.7 | 0 | 0 | 0 | 0.18 | 0.51 | 0.54 | Neiva et al., 2015 |
| Cystoseira amentacea | msat | Fst | | 55 | 55 | -47.82 | -48.83 | -54.32 | 0 | 0 | 0 | 0.26 | 0.28 | 0.35 | Bermejo et al., 2018 |
| Cystoseira amentacea | msat | Fst | | 15 | 15 | -26.3 | -36.67 | -33.07 | 0.03 | 0 | 0 | 0.27 | 0.63 | 0.53 | Buonomo et al., 2017 |
| Cystoseira mediterranea | msat | Fst | | 6 | 6 | NA | NA | NA | NA | NA | NA | NA | NA | NA | Bermejo et al., 2018 |
| Cystoseira tamariscifolia | msat | Fst | | 171 | 171 | -157.74 | -270.3 | -277.76 | 0 | 0 | 0 | 0.35 | 0.67 | 0.68 | Bermejo et al., 2018 |
| Ecklonia radiata | nucDNA | Fst | adaptive | 15 | 15 | -24.05 | -27.4 | -25.65 | 0 | 0 | 0 | 0.57 | 0.65 | 0.61 | Vranken et al., 2021 |
| Ecklonia radiata | nucDNA | Fst | neutral | 15 | 15 | -8.99 | -8.79 | -12.28 | 1 | 0.22 | 0.04 | 0 | 0.05 | 0.24 | Vranken et al., 2021 |
| Ecklonia radiata | msat | Fst | beforevsafter | 10 | 10 | -39.31 | -63.46 | -63.82 | 0.03 | 0 | 0 | 0.39 | 0.95 | 0.95 | Coleman et al., 2020 |
| Ecklonia radiata | msat | Fst | before | 6 | 6 | NA | NA | NA | NA | NA | NA | NA | NA | NA | Coleman et al., 2020 |
| Ecklonia radiata | msat | Fst | after | 6 | 6 | NA | NA | NA | NA | NA | NA | NA | NA | NA | Coleman et al., 2020 |
| Ericaria zosteroides | nucDNA | Fst | | 10 | 10 | -56.18 | -54.23 | -54.27 | 1 | 0.84 | 0.79 | 0 | 0 | 0 | Reynes et al., 2021 |
| Fucus ceranoides | msat | Fst | | 171 | 171 | -131.6 | -354.01 | -365.11 | 1 | 0 | 0 | 0 | 0.73 | 0.75 | Neiva et al., 2012 |
| Fucus ceranoides | msat | JostD | | 171 | 171 | -77.59 | -281.02 | -290.04 | 0 | 0 | 0 | 0.24 | 0.77 | 0.78 | Neiva et al., 2012 |
| Fucus ceranoides | msat | Fst | | 210 | 210 | -94.04 | -120.05 | -119.13 | 0 | 0 | 0 | 0.13 | 0.23 | 0.22 | Neiva et al., 2012 |
| Fucus ceranoides | msat | Dst | | 210 | 210 | -197.5 | -402.15 | -406.41 | 0 | 0 | 0 | 0.13 | 0.67 | 0.68 | Neiva et al., 2012 |
| Fucus distichus | msat | Fst | | 1 | 1 | NA | NA | NA | NA | NA | NA | NA | NA | NA | Coleman et al., 2005 |
| Fucus guiryi | msat | Fst | | 136 | 135 | -98.28 | -98.13 | -101.52 | 0 | 0 | 0 | 0.34 | 0.34 | 0.35 | Lourenço et al., 2016 |
| Fucus guiryi | msat | JostD | | 136 | 135 | -142.64 | -142.75 | -141.25 | 0 | 0 | 0 | 0.22 | 0.22 | 0.22 | Lourenço et al., 2016 |
| Fucus radicans | msat | Fst | | 10 | 10 | -20.92 | -22.28 | -22.89 | 0.02 | 0.01 | 0.01 | 0.42 | 0.5 | 0.53 | Pereyra et al., 2013 |
| Fucus spiralis | msat | Fst | | 1 | 1 | NA | NA | NA | NA | NA | NA | NA | NA | NA | Coleman et al., 2005 |
| Fucus vesiculosus | msat | Fst | | 153 | 153 | -264.75 | -360.16 | -369.5 | 0 | 0 | 0 | 0.38 | 0.67 | 0.69 | Assis et al., 2014 |
| Fucus vesiculosus | msat | JostD | | 153 | 153 | -105.08 | -216.72 | -219.97 | 0 | 0 | 0 | 0.32 | 0.67 | 0.68 | Assis et al., 2014 |
| Fucus vesiculosus | msat | Fst | | 10 | 10 | -20.58 | -21.73 | -23.59 | 1 | 0.12 | 0.05 | 0 | 0.18 | 0.32 | Pereyra et al., 2013 |
| Fucus vesiculosus | msat | Fst | | 55 | 55 | -107.34 | -107.39 | -106.59 | 0 | 0 | 0 | 0.38 | 0.38 | 0.37 | Nicastro et al., 2013 |

| | | | | | | AIC | | | *p*-value | | | $R^2$ | | | |
|---|---|---|---|---|---|---|---|---|---|---|---|---|---|---|---|
| **Species** | **Marker** | **Diff.Index** | **Spec** | **Pairs** | **Pairs.Co** | **SM** | **CM** | **CCM** | **SM** | **CM** | **CCM** | **SM** | **CM** | **CCM** | **Reference** |
| Fucus vesiculosus | msat | Dst | | 55 | 55 | -10.06 | -13.52 | -14.91 | 1 | 0.02 | 0.01 | 0 | 0.08 | 0.1 | Nicastro et al., 2013 |
| Fucus vesiculosus | msat | Fst | | 10 | 10 | -39.8 | -36.67 | -37.23 | 0.01 | 0.03 | 0.02 | 0 | 0.4 | 0.43 | Zardi et al., 2013 |
| Halidrys dioica | isozyme | Nei | | 1 | 1 | NA | NA | NA | NA | NA | NA | NA | NA | NA | Lu et al., 1994 |
| Hormosira banksii | msat | Fst | | 15 | 15 | -48.15 | -49.43 | -49.58 | 0 | 0 | 0 | 0.67 | 0.7 | 0.7 | Coleman et al., 2019 |
| Hormosira banksii | mtDNA | Fst | | 171 | 171 | 74.26 | 73.96 | 70.05 | 0 | 0 | 0 | 0.43 | 0.44 | 0.45 | Mueller et al., 2018 |
| Ishige okamurae | mtDNA | Fst | | 55 | 55 | 4.76 | 9.34 | 16.64 | 0 | 0 | 0 | 0 | 0.38 | 0.29 | Lee et al., 2012 |
| Laminaria digitata | msat | Fst | | 1 | 1 | NA | NA | NA | NA | NA | NA | NA | NA | NA | Brennan et al., 2014 |
| Laminaria digitata | msat | Fst | | 45 | 45 | -120.96 | -187.76 | -195.79 | 0 | 0 | 0 | 0.38 | 0.86 | 0.88 | Robuchon et al., 2014 |
| Laminaria hyperborea | msat | Fst | | 45 | 45 | -93.63 | -164.86 | -205.74 | 1 | 0 | 0 | 0 | 0.8 | 0.92 | Robuchon et al., 2014 |
| Laminaria ochroleuca | msat | JostD | | 171 | 171 | -367.28 | -413.33 | -415.56 | 0 | 0 | 0 | 0.84 | 0.88 | 0.88 | Assis et al., 2018 |
| Laminaria pallida | msat | Fst | | 105 | 105 | -279.29 | -475.28 | -477.56 | 0 | 0 | 0 | 0.38 | 0.9 | 0.91 | Assis et al., 2022 |
| Macrocystis pyrifera | msat | Fst | Southern hemisphere | 666 | 595 | -789.84 | -793.34 | -794.03 | 0 | 0 | 0 | 0.25 | 0.25 | 0.26 | Assis et al., 2023 |
| Macrocystis pyrifera | msat | Fst | Northern hemisphere | 528 | 528 | -731.91 | -756.54 | -762.62 | 0 | 0 | 0 | 0.26 | 0.29 | 0.3 | Assis et al., 2023 |
| Nereia lophocladia | nucDNA | Fst | | 6 | 6 | NA | NA | NA | NA | NA | NA | NA | NA | NA | Mamo et al., 2021 |
| Phyllospora comosa | nucDNA | Fst | | 66 | 66 | -148.56 | -206.61 | -216.23 | 0 | 0 | 0 | 0.69 | 0.87 | 0.89 | Wood et al., 2021 |
| Pylaiella littoralis | msat | Fst | | 6 | 6 | NA | NA | NA | NA | NA | NA | NA | NA | NA | Geoffroy et al., 2015 |
| Saccharina japonica | msat | Fst | | 465 | 465 | -851.33 | -949.64 | -951.16 | 0 | 0 | 0 | 0.35 | 0.47 | 0.47 | Zhang et al., 2019 |
| Saccharina japonica | mtDNA | Fst | | 465 | 465 | 43.23 | 34.58 | -59.59 | 0 | 0 | 0 | 0.08 | 0.1 | 0.26 | Zhang et al., 2019 |
| Saccharina japonica | mtDNA | Fst | | 210 | 210 | -33.55 | -35.73 | -35.84 | 0 | 0 | 0 | 0.63 | 0.63 | 0.63 | Zhang et al., 2015 |
| Saccharina japonica | nucDNA | Fst | | 1 | 1 | NA | NA | NA | NA | NA | NA | NA | NA | NA | Zhao et al., 2013 |
| Saccharina japonica | nucDNA | Nei | | 1 | 1 | NA | NA | NA | NA | NA | NA | NA | NA | NA | Zhao et al., 2013 |
| Saccharina latissima | msat | Fst | | 28 | 28 | -69.06 | -101.79 | -97.73 | 1 | 0 | 0 | 0 | 0.7 | 0.65 | Møller Nielsen et al., 2016 |

| | | | | | | AIC | | | *p*-value | | | $R^2$ | | | |
|---|---|---|---|---|---|---|---|---|---|---|---|---|---|---|---|
| **Species** | **Marker** | **Diff.Index** | **Spec** | **Pairs** | **Pairs.Co** | **SM** | **CM** | **CCM** | **SM** | **CM** | **CCM** | **SM** | **CM** | **CCM** | **Reference** |
| Saccharina latissima | msat | Fst | | 6 | 6 | NA | NA | NA | NA | NA | NA | NA | NA | NA | Breton et al., 2018 |
| Saccharina latissima | nucDNA | Fst | | 55 | 55 | -52.17 | -93.33 | -100.73 | 0 | 0 | 0 | 0.29 | 0.66 | 0.71 | Guzinski et al., 2020 |
| Saccorhiza polyschides | msat | Fst | | 351 | 351 | -530.82 | -842.09 | -843.92 | 0 | 0 | 0 | 0.19 | 0.67 | 0.67 | Assis et al., 2016 |
| Saccorhiza polyschides | msat | JostD | | 351 | 351 | -344.2 | -491.63 | -492.22 | 0 | 0 | 0 | 0.46 | 0.64 | 0.64 | Assis et al., 2016 |
| Saccorhiza polyschides | msat | Fst | | 91 | 91 | -292.96 | -330.5 | -336.34 | 0 | 0 | 0 | 0.74 | 0.83 | 0.84 | Assis et al., 2013 |
| Saccorhiza polyschides | msat | JostD | | 91 | 91 | -207.93 | -260.46 | -266.46 | 0 | 0 | 0 | 0.78 | 0.88 | 0.88 | Assis et al., 2013 |
| Sargassum fallax | mtDNA | Gst | before | 2 | 2 | NA | NA | NA | NA | NA | NA | NA | NA | NA | Gurgel et al., 2020 |
| Sargassum fallax | mtDNA | Gst | after | 3 | 3 | NA | NA | NA | NA | NA | NA | NA | NA | NA | Gurgel et al., 2020 |
| Sargassum fallax | cpDNA | Gst | before | 6 | 6 | NA | NA | NA | NA | NA | NA | NA | NA | NA | Gurgel et al., 2020 |
| Sargassum fallax | cpDNA | Gst | after | 10 | 10 | -6.7 | -5.96 | -6.27 | 1 | 0.33 | 0.28 | 0 | 0.01 | 0.04 | Gurgel et al., 2020 |
| Sargassum fusiforme | mtDNA | Fst | | 300 | 300 | 181.7 | 56.85 | 42.09 | 0 | 0 | 0 | 0.11 | 0.41 | 0.44 | Hu et al., 2017 |
| Sargassum fusiforme | cpDNA | Fst | | 300 | 300 | -187.56 | -186.88 | -186.89 | 1 | 0.25 | 0.25 | 0 | 0 | 0 | Hu et al., 2017 |
| Sargassum fusiforme | mtDNA | Fst | | 45 | 45 | 22.18 | -25.69 | -49.06 | 0 | 0 | 0 | 0.22 | 0.73 | 0.84 | Hu et al., 2013 |
| Sargassum fusiforme | nucDNA | Fst | | 36 | 36 | -148.35 | -149.1 | -161.13 | 0 | 0 | 0 | 0.66 | 0.67 | 0.76 | Yu et al., 2012 |
| Sargassum fusiforme | nucDNA | Nei | | 36 | 36 | -151.17 | -166.95 | -166.98 | 0 | 0 | 0 | 0.28 | 0.54 | 0.54 | Yu et al., 2012 |
| Sargassum hemiphyllum | nucDNA | Fst | | 36 | 36 | 30.56 | 2.09 | -4.55 | 0 | 0 | 0 | 0.52 | 0.78 | 0.82 | Cheang et al., 2010 |
| Sargassum hemiphyllum | cpDNA | Fst | | 36 | 36 | 31.36 | 2.27 | -5.74 | 0 | 0 | 0 | 0.37 | 0.72 | 0.78 | Cheang et al., 2010 |
| Sargassum hemiphyllum | mtDNA | Fst | | 36 | 36 | 14.18 | -4.57 | -6.23 | 0 | 0 | 0 | 0.34 | 0.61 | 0.62 | Cheang et al., 2010 |
| Sargassum horneri | mtDNA | Gst | | 78 | 78 | 50.49 | 45.36 | 45.53 | 0 | 0 | 0 | 0.18 | 0.24 | 0.23 | Watanabe et al., 2019 |
| Sargassum horneri | mtDNA | Nst | | 78 | 78 | 66.26 | 60.28 | 54.39 | 1 | 0.01 | 0 | 0 | 0.09 | 0.15 | Watanabe et al., 2019 |
| Sargassum horneri | nucDNA | Nei | | 36 | 36 | -92.88 | -91.03 | -93.76 | 1 | 0.71 | 0.1 | 0 | 0 | 0.05 | Yu et al., 2013 |

| | | | | | | AIC | | | p-value | | | $R^2$ | | | |
|---|---|---|---|---|---|---|---|---|---|---|---|---|---|---|---|
| **Species** | **Marker** | **Diff.Index** | **Spec** | **Pairs** | **Pairs.Co** | **SM** | **CM** | **CCM** | **SM** | **CM** | **CCM** | **SM** | **CM** | **CCM** | **Reference** |
| Sargassum ilicifolium | nucDNA | Fst | | 105 | 105 | -27.41 | -17.58 | -18.1 | 0 | 0 | 0 | 0 | 0.13 | 0.14 | Ng et al., 2019 |
| Sargassum ilicifolium | mtDNA | Fst | | 105 | 105 | -28.52 | -28.08 | -20.65 | 0 | 0 | 0 | 0 | 0.2 | 0.14 | Ng et al., 2019 |
| Sargassum polycystum | mtDNA | PHIst | | 10 | 10 | -10.39 | -9.78 | -14.1 | 0 | 0 | 0 | 0.88 | 0.87 | 0.91 | Kanyachumpoo et al., 2014 |
| Sargassum polycystum | nucDNA | PHIst | | 325 | 325 | 308.85 | 301.04 | 300.31 | 0 | 0 | 0 | 0.25 | 0.26 | 0.27 | Chan et al., 2013 |
| Sargassum polycystum | cpDNA | PHIst | | 190 | 190 | 79.85 | 99.21 | 99.07 | 0 | 0 | 0 | 0 | 0.06 | 0.06 | Chan et al., 2013 |
| Sargassum polycystum | mtDNA | PHIst | | 153 | 153 | 48.97 | 45.03 | 43.63 | 0 | 0 | 0 | 0.59 | 0.6 | 0.61 | Chan et al., 2013 |
| Sargassum thunbergii | nucDNA | Fst | | 406 | 406 | -326.81 | -338.58 | -354.28 | 0 | 0 | 0 | 0.39 | 0.41 | 0.43 | Li et al., 2017 |
| Sargassum thunbergii | mtDNA | Fst | | 406 | 406 | -35.4 | -46.17 | -71.03 | 0 | 0 | 0 | 0.18 | 0.2 | 0.25 | Li et al., 2017 |
| Sargassum thunbergii | msat | Fst | | 378 | 378 | -498.31 | -591.61 | -626.08 | 0 | 0 | 0 | 0.18 | 0.36 | 0.41 | Song et al., 2021 |
| Sargassum thunbergii | msat | JostD | | 378 | 378 | -221.68 | -302.15 | -372.61 | 0 | 0 | 0 | 0.09 | 0.26 | 0.39 | Song et al., 2021 |
| Undaria pinnatifida | msat | Fst | native | 3 | 3 | NA | NA | NA | NA | NA | NA | NA | NA | NA | Shan et al., 2019 |
| Undaria pinnatifida | msat | Fst | invasive | 15 | 15 | -16.86 | -30.1 | -32.59 | 0.04 | 0 | 0 | 0.24 | 0.69 | 0.73 | Shan et al., 2019 |
| Undaria pinnatifida | msat | Nei | native | 3 | 3 | NA | NA | NA | NA | NA | NA | NA | NA | NA | Shan et al., 2019 |
| Undaria pinnatifida | msat | Nei | invasive | 15 | 15 | 1.56 | -0.74 | -2.16 | 1 | 0.06 | 0.03 | 0 | 0.19 | 0.26 | Shan et al., 2019 |
| Undaria pinnatifida | nucDNA | Fst | | 66 | 66 | -149.99 | -150.78 | -151.33 | 0 | 0 | 0 | 0.41 | 0.41 | 0.42 | Guzinski et al., 2018 |
| Undaria pinnatifida | msat | Fst | | 66 | 66 | -144.13 | -144.87 | -144.73 | 0 | 0 | 0 | 0.24 | 0.25 | 0.25 | Guzinski et al., 2018 |

Table S4: Performance of Spatial Model (SM), Connectivity Model (CM) and Connectivity and Centrality Model (CCM), using a fixed PD value (i.e. 7.43 d). Species, genetic marker, genetic differentiation index (Diff.Index), dataset speciality (Spec), number of pairs of sites (Pairs), number of pairs of sites connected (Pairs.Co) and dataset reference are reported. Performance of models are given with Akaike information criterion (AIC), significance levels (p-value) and the goodness of fit ($R^2$).

| | | | | | | **AIC** | | | ***p*-value** | | | **R²** | | | |
|---|---|---|---|---|---|---|---|---|---|---|---|---|---|---|---|
| **Species** | **Marker** | **Diff.Index** | **Spec** | **Pairs** | **Pairs.Co** | **SM** | **CM** | **CCM** | **SM** | **CM** | **CCM** | **SM** | **CM** | **CCM** | **Reference** |
| Bifurcaria bifurcata | msat | Fst | | 153 | 56 | -140.85 | -155.21 | -155.18 | 0 | 0 | 0 | 0.39 | 0.44 | 0.44 | Neiva et al., 2015 |
| Bifurcaria bifurcata | msat | JostD | | 153 | 56 | -244.54 | -292.7 | -317.63 | 0 | 0 | 0 | 0.18 | 0.4 | 0.49 | Neiva et al., 2015 |
| Cystoseira amentacea | msat | Fst | | 55 | 49 | -47.82 | -47.59 | -39.25 | 0 | 0 | 0 | 0.26 | 0.26 | 0.14 | Bermejo et al., 2018 |
| Cystoseira amentacea | msat | Fst | | 15 | 15 | -26.3 | -35.18 | -33.32 | 0.03 | 0 | 0 | 0.27 | 0.6 | 0.54 | Buonomo et al., 2017 |
| Cystoseira mediterranea | msat | Fst | | 6 | 3 | NA | NA | NA | NA | NA | NA | NA | NA | NA | Bermejo et al., 2018 |
| Cystoseira tamariscifolia | msat | Fst | | 171 | 73 | -157.74 | -205.26 | -228.83 | 0 | 0 | 0 | 0.35 | 0.51 | 0.57 | Bermejo et al., 2018 |
| Ecklonia radiata | nucDNA | Fst | adaptive | 15 | 15 | -24.05 | -45.03 | -45.06 | 0 | 0 | 0 | 0.57 | 0.89 | 0.89 | Vranken et al., 2021 |
| Ecklonia radiata | nucDNA | Fst | neutral | 15 | 15 | -8.99 | -14.59 | -15.1 | NA | 0.01 | 0.01 | 0 | 0.35 | 0.37 | Vranken et al., 2021 |
| Ecklonia radiata | msat | Fst | beforevsafter | 10 | 10 | -39.31 | -49.44 | -55.49 | 0.03 | 0 | 0 | 0.39 | 0.78 | 0.88 | Coleman et al., 2020 |
| Ecklonia radiata | msat | Fst | before | 6 | 6 | NA | NA | NA | NA | NA | NA | NA | NA | NA | Coleman et al., 2020 |
| Ecklonia radiata | msat | Fst | after | 6 | 6 | NA | NA | NA | NA | NA | NA | NA | NA | NA | Coleman et al., 2020 |
| Ericaria zosteroides | nucDNA | Fst | | 10 | 6 | -56.18 | -54.6 | -58.23 | NA | 0.57 | 0.08 | 0 | 0 | 0.25 | Reynes et al., 2021 |
| Fucus ceranoides | msat | Fst | | 171 | 171 | -131.6 | -355.71 | -364.15 | NA | 0 | 0 | 0 | 0.73 | 0.74 | Neiva et al., 2012 |
| Fucus ceranoides | msat | JostD | | 171 | 171 | -77.59 | -307.66 | -313.29 | 0 | 0 | 0 | 0.24 | 0.8 | 0.81 | Neiva et al., 2012 |
| Fucus ceranoides | msat | Fst | | 210 | 49 | -94.04 | -84.44 | -83.84 | 0 | 0 | 0 | 0.13 | 0.09 | 0.08 | Neiva et al., 2012 |
| Fucus ceranoides | msat | Dst | | 210 | 49 | -197.5 | -171.94 | -175.73 | 0 | 0.05 | 0.01 | 0.13 | 0.01 | 0.03 | Neiva et al., 2012 |
| Fucus distichus | msat | Fst | | 1 | 1 | NA | NA | NA | NA | NA | NA | NA | NA | NA | Coleman et al., 2005 |
| Fucus guiryi | msat | Fst | | 136 | 77 | -98.28 | -105.15 | -107.1 | 0 | 0 | 0 | 0.34 | 0.37 | 0.38 | Lourenço et al., 2016 |
| Fucus guiryi | msat | JostD | | 136 | 77 | -142.64 | -142.17 | -144.75 | 0 | 0 | 0 | 0.22 | 0.22 | 0.24 | Lourenço et al., 2016 |
| Fucus radicans | msat | Fst | | 10 | 2 | -20.92 | -21 | -20.56 | 0.02 | 0.02 | 0.03 | 0.42 | 0.43 | 0.4 | Pereyra et al., 2013 |
| Fucus spiralis | msat | Fst | | 1 | 1 | NA | NA | NA | NA | NA | NA | NA | NA | NA | Coleman et al., 2005 |
| Fucus vesiculosus | msat | Fst | | 153 | 38 | -264.75 | -266.27 | -269.6 | 0 | 0 | 0 | 0.38 | 0.38 | 0.4 | Assis et al., 2014 |
| Fucus vesiculosus | msat | JostD | | 153 | 38 | -105.08 | -114.75 | -117.84 | 0 | 0 | 0 | 0.32 | 0.36 | 0.37 | Assis et al., 2014 |
| Fucus vesiculosus | msat | Fst | | 10 | 2 | -20.58 | -24.75 | -25.24 | NA | 0.03 | 0.03 | 0 | 0.39 | 0.42 | Pereyra et al., 2013 |
| Fucus vesiculosus | msat | Fst | | 55 | 45 | -107.34 | -104.06 | -112.72 | 0 | 0 | 0 | 0.38 | 0.34 | 0.44 | Nicastro et al., 2013 |

| | | | | | | AIC | | | *p*-value | | | $R^2$ | | | |
|---|---|---|---|---|---|---|---|---|---|---|---|---|---|---|---|
| **Species** | **Marker** | **Diff.Index** | **Spec** | **Pairs** | **Pairs.Co** | **SM** | **CM** | **CCM** | **SM** | **CM** | **CCM** | **SM** | **CM** | **CCM** | **Reference** |
| Fucus vesiculosus | msat | Dst | | 55 | 45 | -10.06 | -8.78 | -8.78 | NA | 0.41 | 0.41 | 0 | 0 | 0 | Nicastro et al., 2013 |
| Fucus vesiculosus | msat | Fst | | 10 | 4 | -39.8 | -36.83 | -37.1 | 0.01 | 0.03 | 0.02 | 0.56 | 0.41 | 0.42 | Zardi et al., 2013 |
| Halidrys dioica | isozyme | Nei | | 1 | 1 | NA | NA | NA | NA | NA | NA | NA | NA | NA | Coleman et al., 2019 |
| Hormosira banksii | msat | Fst | | 15 | 15 | -48.15 | -49.5 | -50.02 | 0 | 0 | 0 | 0.67 | 0.7 | 0.71 | Mueller et al., 2018 |
| Hormosira banksii | mtDNA | Fst | | 171 | 76 | 74.26 | 71.76 | 62.01 | 0 | 0 | 0 | 0.43 | 0.44 | 0.47 | Lu et al., 1994 |
| Ishige okamurae | mtDNA | Fst | | 55 | 7 | 4.76 | 3.93 | 4.76 | 0 | 0 | 0 | 0.43 | 0.43 | 0.43 | Lee et al., 2012 |
| Laminaria digitata | msat | Fst | | 1 | 1 | NA | NA | NA | NA | NA | NA | NA | NA | NA | Brennan et al., 2014 |
| Laminaria digitata | msat | Fst | | 45 | 45 | -120.96 | -203.81 | -241.73 | 0 | 0 | 0 | 0.38 | 0.9 | 0.96 | Robuchon et al., 2014 |
| Laminaria hyperborea | msat | Fst | | 45 | 45 | -93.63 | -156.16 | -200.71 | NA | 0 | 0 | 0 | 0.76 | 0.91 | Robuchon et al., 2014 |
| Laminaria ochroleuca | msat | JostD | | 171 | 105 | -367.28 | -388.41 | -391.8 | 0 | 0 | 0 | 0.84 | 0.86 | 0.86 | Assis et al., 2018 |
| Laminaria pallida | msat | Fst | | 105 | 105 | -279.29 | -461.13 | -462.22 | 0 | 0 | 0 | 0.38 | 0.89 | 0.89 | Assis et al., 2022 |
| Macrocystis pyrifera | msat | Fst | Southern hemisphere | 666 | 139 | -789.84 | -811.92 | -814.17 | 0 | 0 | 0 | 0.25 | 0.28 | 0.28 | Assis et al., 2023 |
| Macrocystis pyrifera | msat | Fst | Northern hemisphere | 528 | 406 | -731.91 | -756.67 | -962.04 | 0 | 0 | 0 | 0.26 | 0.29 | 0.52 | Assis et al., 2023 |
| Nereia lophocladia | nucDNA | Fst | | 6 | 6 | NA | NA | NA | NA | NA | NA | NA | NA | NA | Mamo et al., 2021 |
| Phyllospora comosa | nucDNA | Fst | | 66 | 46 | -148.56 | -151.87 | -153.55 | 0 | 0 | 0 | 0.69 | 0.71 | 0.72 | Wood et al., 2021 |
| Pylaiella littoralis | msat | Fst | | 6 | 6 | NA | NA | NA | NA | NA | NA | NA | NA | NA | Geoffroy et al., 2015 |
| Saccharina japonica | msat | Fst | | 465 | 259 | -851.33 | -883.69 | -889.51 | 0 | 0 | 0 | 0.35 | 0.39 | 0.4 | Gurgel et al., 2020 |
| Saccharina japonica | mtDNA | Fst | | 465 | 259 | 43.23 | -23.34 | -46.45 | 0 | 0 | 0 | 0.08 | 0.2 | 0.24 | Gurgel et al., 2020 |
| Saccharina japonica | mtDNA | Fst | | 210 | 109 | -33.55 | -32.97 | -34.3 | 0 | 0 | 0 | 0.63 | 0.63 | 0.63 | Gurgel et al., 2020 |
| Saccharina japonica | nucDNA | Fst | | 1 | 1 | NA | NA | NA | NA | NA | NA | NA | NA | NA | Gurgel et al., 2020 |
| Saccharina japonica | nucDNA | Nei | | 1 | 1 | NA | NA | NA | NA | NA | NA | NA | NA | NA | Hu et al., 2017 |
| Saccharina latissima | msat | Fst | | 28 | 28 | -69.06 | -81.59 | -84.92 | NA | 0 | 0 | 0 | 0.38 | 0.45 | Hu et al., 2017 |

| | | | | | | AIC | | | *p*-value | | | $R^2$ | | | |
|---|---|---|---|---|---|---|---|---|---|---|---|---|---|---|---|
| **Species** | **Marker** | **Diff.Index** | **Spec** | **Pairs** | **Pairs.Co** | **SM** | **CM** | **CCM** | **SM** | **CM** | **CCM** | **SM** | **CM** | **CCM** | **Reference** |
| Saccharina latissima | msat | Fst | | 6 | 6 | NA | NA | NA | NA | NA | NA | NA | NA | NA | Hu et al., 2013 |
| Saccharina latissima | nucDNA | Fst | | 55 | 18 | -52.17 | -49.22 | -50.39 | 0 | 0 | 0 | 0.29 | 0.25 | 0.26 | Yu et al., 2012 |
| Saccorhiza polyschides | msat | Fst | | 351 | 190 | -530.82 | -677.55 | -677.68 | 0 | 0 | 0 | 0.19 | 0.47 | 0.47 | Yu et al., 2012 |
| Saccorhiza polyschides | msat | JostD | | 351 | 190 | -344.2 | -431.13 | -438.91 | 0 | 0 | 0 | 0.46 | 0.58 | 0.58 | Cheang et al., 2010 |
| Saccorhiza polyschides | msat | Fst | | 91 | 91 | -292.96 | -324.08 | -324.2 | 0 | 0 | 0 | 0.74 | 0.81 | 0.81 | Cheang et al., 2010 |
| Saccorhiza polyschides | msat | JostD | | 91 | 91 | -207.93 | -252.91 | -253.15 | 0 | 0 | 0 | 0.78 | 0.87 | 0.87 | Cheang et al., 2010 |
| Sargassum fallax | mtDNA | Gst | before | 2 | 2 | NA | NA | NA | NA | NA | NA | NA | NA | NA | Watanabe et al., 2019 |
| Sargassum fallax | mtDNA | Gst | after | 3 | 3 | NA | NA | NA | NA | NA | NA | NA | NA | NA | Watanabe et al., 2019 |
| Sargassum fallax | cpDNA | Gst | before | 6 | 6 | NA | NA | NA | NA | NA | NA | NA | NA | NA | Yu et al., 2013 |
| Sargassum fallax | cpDNA | Gst | after | 10 | 10 | -6.7 | -6.61 | -6.79 | NA | 0.23 | 0.21 | 0 | 0.07 | 0.09 | Ng et al., 2019 |
| Sargassum fusiforme | mtDNA | Fst | | 300 | 85 | 181.7 | 109.8 | 90.34 | 0 | 0 | 0 | 0.11 | 0.3 | 0.34 | Ng et al., 2019 |
| Sargassum fusiforme | cpDNA | Fst | | 300 | 85 | -187.56 | -186.02 | -190.37 | NA | 0.5 | 0.03 | 0 | 0 | 0.01 | Zhang et al., 2019 |
| Sargassum fusiforme | mtDNA | Fst | | 45 | 27 | 22.18 | 18.19 | 14.76 | 0 | 0 | 0 | 0.22 | 0.29 | 0.34 | Zhang et al., 2019 |
| Sargassum fusiforme | nucDNA | Fst | | 36 | 27 | -148.35 | -157.13 | -157.46 | 0 | 0 | 0 | 0.66 | 0.73 | 0.73 | Zhang et al., 2015 |
| Sargassum fusiforme | nucDNA | Nei | | 36 | 27 | -151.17 | -167.62 | -167.92 | 0 | 0 | 0 | 0.28 | 0.55 | 0.55 | Zhao et al., 2013 |
| Sargassum hemiphyllum | nucDNA | Fst | | 36 | 9 | 30.56 | 14.12 | 11.78 | 0 | 0 | 0 | 0.52 | 0.69 | 0.71 | Zhao et al., 2013 |
| Sargassum hemiphyllum | cpDNA | Fst | | 36 | 9 | 31.36 | 16.07 | 15.85 | 0 | 0 | 0 | 0.37 | 0.59 | 0.59 | Møller Nielsen et al., 2016 |
| Sargassum hemiphyllum | mtDNA | Fst | | 36 | 9 | 14.18 | 7.02 | 4.83 | 0 | 0 | 0 | 0.34 | 0.46 | 0.49 | Breton et al., 2018 |
| Sargassum horneri | mtDNA | Gst | | 78 | 66 | 50.49 | 42.03 | 42.72 | 0 | 0 | 0 | 0.18 | 0.27 | 0.26 | Guzinski et al., 2020 |
| Sargassum horneri | mtDNA | Nst | | 78 | 66 | 66.26 | 47.55 | 47.25 | NA | 0 | 0 | 0 | 0.22 | 0.23 | Kanyachumpoo et al., 2014 |

| | | | | | | AIC | | | *p*-value | | | $R^2$ | | | |
|---|---|---|---|---|---|---|---|---|---|---|---|---|---|---|---|
| **Species** | **Marker** | **Diff.Index** | **Spec** | **Pairs** | **Pairs.Co** | **SM** | **CM** | **CCM** | **SM** | **CM** | **CCM** | **SM** | **CM** | **CCM** | **Reference** |
| Sargassum horneri | nucDNA | Nei | | 36 | 25 | -92.88 | -91.73 | -94.45 | NA | 0.38 | 0.07 | 0 | 0 | 0.07 | Chan et al., 2013 |
| Sargassum ilicifolium | nucDNA | Fst | | 105 | 49 | -27.41 | -12.19 | -11.6 | 0 | 0 | 0 | 0.21 | 0.09 | 0.08 | Chan et al., 2013 |
| Sargassum ilicifolium | mtDNA | Fst | | 105 | 49 | -28.52 | -36.15 | -33.79 | 0 | 0 | 0 | 0.2 | 0.26 | 0.24 | Chan et al., 2013 |
| Sargassum polycystum | mtDNA | PHIst | | 10 | 1 | -10.39 | -10.7 | -17.54 | 0 | 0 | 0 | 0.88 | 0.88 | 0.94 | Assis et al., 2016 |
| Sargassum polycystum | nucDNA | PHIst | | 325 | 26 | 308.85 | 308.03 | 306.26 | 0 | 0 | 0 | 0.25 | 0.25 | 0.25 | Assis et al., 2016 |
| Sargassum polycystum | cpDNA | PHIst | | 190 | 17 | 79.85 | 78.69 | 84.05 | 0 | 0 | 0 | 0.15 | 0.16 | 0.13 | Assis et al., 2013 |
| Sargassum polycystum | mtDNA | PHIst | | 153 | 9 | 48.97 | 48.04 | 47.73 | 0 | 0 | 0 | 0.59 | 0.59 | 0.6 | Assis et al., 2013 |
| Sargassum thunbergii | nucDNA | Fst | | 406 | 119 | -326.81 | -331.46 | -337.34 | 0 | 0 | 0 | 0.39 | 0.4 | 0.41 | Li et al., 2017 |
| Sargassum thunbergii | mtDNA | Fst | | 406 | 119 | -35.4 | -55.57 | -49.68 | 0 | 0 | 0 | 0.18 | 0.22 | 0.2 | Li et al., 2017 |
| Sargassum thunbergii | msat | Fst | | 378 | 117 | -498.31 | -576.86 | -667.16 | 0 | 0 | 0 | 0.18 | 0.33 | 0.47 | Song et al., 2021 |
| Sargassum thunbergii | msat | JostD | | 378 | 117 | -221.68 | -362.83 | -436.76 | 0 | 0 | 0 | 0.09 | 0.37 | 0.48 | Song et al., 2021 |
| Undaria pinnatifida | msat | Fst | native | 3 | 1 | NA | NA | NA | NA | NA | NA | NA | NA | NA | Shan et al., 2019 |
| Undaria pinnatifida | msat | Fst | invasive | 15 | 4 | -16.86 | -17.15 | -48.83 | 0.04 | 0.03 | 0 | 0.24 | 0.26 | 0.91 | Shan et al., 2019 |
| Undaria pinnatifida | msat | Nei | native | 3 | 1 | NA | NA | NA | NA | NA | NA | NA | NA | NA | Shan et al., 2019 |
| Undaria pinnatifida | msat | Nei | invasive | 15 | 4 | 1.56 | 1.29 | 1.28 | NA | 0.17 | 0.17 | 0 | 0.07 | 0.07 | Shan et al., 2019 |
| Undaria pinnatifida | nucDNA | Fst | | 66 | 66 | -149.99 | -150.25 | -151.9 | 0 | 0 | 0 | 0.41 | 0.41 | 0.42 | Guzinski et al., 2018 |
| Undaria pinnatifida | msat | Fst | | 66 | 66 | -144.13 | -144.47 | -142.63 | 0 | 0 | 0 | 0.24 | 0.24 | 0.22 | Guzinski et al., 2018 |

Table S5: Sensitivity of the performance of Spatial Model (SM) and Connectivity and Centrality Model (CCM) on dataset characteristics. The characteristics displayed are the genetic marker (Marker), the genetic differentiation index (Diff.Index), the species phylogeny (Genus, Family, Order), the sampled realm, the number of populations (n), the maximum distance between sampling sites (Samp.Ext), the mean distance between sampling sites (Samp.Mean.Dist), and the latitude extent between sampling sites (Lat.Ext).

| **Model** | | **Characteristic** | **Statistical test** | ***p*-value** | **$R^2$** |
|---|---|---|---|---|---|
| SM | $R^2$ | Marker | Kruskal-Wallis | 0.57 | NA |
| | | Diff.Index | Kruskal-Wallis | 0.18 | NA |
| | | Genus | Kruskal-Wallis | 0.5 | NA |
| | | Family | Kruskal-Wallis | 0.23 | NA |
| | | Order | Kruskal-Wallis | 0.26 | NA |
| | | Realm | Kruskal-Wallis | 0.84 | NA |
| | | n | Linear model | 0.1 | 0.03 |
| | | Samp.Ext | Linear model | 0.89 | -0.02 |
| | | Samp.Mean.Dist | Linear model | 0.64 | -0.01 |
| | | Lat.Ext | Linear model | 0.79 | -0.02 |
| CCM | $R^2$ | Marker | Kruskal-Wallis | 0.51 | NA |
| | | Diff.Index | Kruskal-Wallis | 0.39 | NA |
| | | Genus | Kruskal-Wallis | 0.09 | NA |
| | | Family | Kruskal-Wallis | 0.25 | NA |
| | | Order | Kruskal-Wallis | 0.16 | NA |
| | | Realm | Kruskal-Wallis | 0.4 | NA |
| | | **n** | **Linear model** | **0.01** | **0.09** |
| | | Samp.Ext | Linear model | 0.06 | 0.04 |
| | | Samp.Mean.Dist | Linear model | 0.19 | 0.01 |
| | | Lat.Ext | Linear model | 0.3 | 0 |
| CCM | $\Delta R^2$ | Marker | Kruskal-Wallis | 0.26 | NA |
| | | Diff.Index | Kruskal-Wallis | 0.4 | NA |
| | | Genus | Kruskal-Wallis | 0.47 | NA |
| | | Family | Kruskal-Wallis | 0.57 | NA |
| | | Order | Kruskal-Wallis | 0.23 | NA |
| | | Realm | Kruskal-Wallis | 0.16 | NA |
| | | n | Linear model | 0.23 | 0.01 |
| | | Samp.Ext | Linear model | 0.05 | 0.04 |
| | | Samp.Mean.Dist | Linear model | 0.06 | 0.04 |
| | | Lat.Ext | Linear model | 0.17 | 0.02 |
